# An optogenetic switch for the Set2 methyltransferase provides evidence for rapid transcription-dependent and independent dynamics of H3K36 methylation

**DOI:** 10.1101/2020.03.28.013706

**Authors:** Andrew M. Lerner, Austin J. Hepperla, Gregory R. Keele, Hashem Meriesh, Hayretin Yumerefendi, David Restrepo, Seth Zimmerman, James Bear, Brian Kuhlman, Ian J. Davis, Brian D. Strahl

## Abstract

Histone H3 lysine 36 methylation (H3K36me) is a conserved histone modification associated with transcription and DNA repair. Although the effects of H3K36 methylation have been studied, the short-term genome-wide dynamics of H3K36me deposition and removal are not known. We established rapid and reversible optogenetic control for Set2, the sole H3K36 methyltransferase in yeast, by fusing the enzyme with the light activated nuclear shuttle (LANS) domain. Early H3K36me3 dynamics identified rapid methylation *in vivo*, with total H3K36me3 levels correlating with RNA abundance. Although genes exhibited disparate levels of H3K36 methylation, relative rates of H3K36me3 accumulation were largely linear and consistent across genes, suggesting a rate-limiting mechanism for H3K36me3 deposition. Removal H3K36me2/3 was also rapid and highly dependent on the demethylase Rph1. However, the per-gene rate of H3K36me3 loss weakly correlated with RNA abundance and followed exponential decay, suggesting H3K36 demethylases act in a global, stochastic manner.

## INTRODUCTION

Histone post-translational modifications (PTMs) are fundamentally involved in both chromatin packaging and in gene regulation (Strahl and Allis 2000; Venkatesh and Workman 2015). Addition and removal of PTMs must be carefully choreographed to regulate engagement of chromatin remodelling complexes and grant cellular machinery access to DNA for transcription, replication, recombination, and DNA repair (Kouzarides 2007). In particular, dynamic regulation of histone methylation and demethylation has been implicated in these processes, ultimately controlling cell fate and differentiation (Greer and Shi 2012; Klose and Zhang 2007).

Histone H3 lysine 36 methylation (H3K36me) is present across eukaryotic organisms and is generated by the methyltransferase Set2 (McDaniel and Strahl 2017; Wagner and Carpenter 2012). As the sole H3K36 methyltransferase in *Saccharomyces cerevisiae* (*S. cerevisiae*), Set2 co-transcriptionally deposits up to three methyl groups, resulting in mono-, di-, or tri-methylated H3K36 (H3K36me1, H3K36me2, and H3K36me3, respectively) (Govind et al. 2010; Keogh et al. 2005; Kizer et al. 2005; Li et al. 2003; Strahl et al. 2002; Xiao 2003). These modifications regulate chromatin structure through diverse pathways, including activation of the histone deacetylase complex Rpd3S (Carrozza et al. 2005; Govind et al. 2010; Joshi and Struhl 2005; Keogh et al. 2005) regulation of histone exchange and transcription elongation (Gilbert et al. 2014; Maltby et al. 2012; Smolle et al. 2012; Venkatesh et al. 2012). These pathways prevent aberrant transcription from cryptic promoters and maintain transcriptional fidelity and genomic stability (Churchman and Weissman 2011; Jha and Strahl 2014; Kim et al. 2016; McDaniel et al. 2017; Neil et al. 2009; Sen et al. 2015). In addition to a role in transcription, H3K36 methylation has also been linked to DNA damage repair, splicing, and cell cycle regulation (McDaniel and Strahl 2017).

H3K36me is primarily removed by two Jumonji domain-containing histone demethylases, Rph1 and Jhd1, which target H3K36me3 and H3K36me2 (Kim and Buratowski 2007; Klose et al. 2007; Tsukada et al. 2006; Tu et al. 2007). Intriguingly, Rph1 was first identified as a damage-responsive repressor of the DNA photolyase *PHR1* (Jang et al. 1999; Kim 2002; Liang et al. 2011), but later as a H3K36 demethylase. Rph1 and Jhd1 exhibit demethylase activity *in vitro* and have been linked to demethylation of H3K36 during transcription elongation *in vivo* (Chang et al. 2011; Kwon and Ahn 2011). Although these demethylases are presumed to function during transcription elongation and in opposition of Set2-dependent H3K36 methylation, recent studies point to at least the ability of Rph1 to function in a non-transcriptional manner to regulate the balance of H3K36me2/3 for processes involved in metabolism and amino acid biosynthesis (Ye et al. 2017). Thus, further investigation of H3K36 demethylation is warranted.

A limitation of studies into H3K36 methylation and demethylation has been the lack of an experimental strategy that is both reversible and that matches the rapid kinetics of deposition and removal of H3K36 methylation. However, new approaches in optogenetics – the use of genetically-encoded, light-responsive proteins to regulate biological processes – offer the minimal latency and rapid reversibility needed to study chromatin state changes (Kim et al. 2017; Liu and Tucker 2017). Several enabling optogenetic tools based on the LOV2 domain of *Avena sativa* phototropin 1 (AsLOV2) have been developed to date (Di Ventura and Kuhlman 2016; Niopek et al. 2014, 2016; Yumerefendi et al. 2015, 2016). Previously, we engineered LOV2 to control protein translocation into and out of the nucleus using light. We used the light activated nuclear shuttle (LANS) to control cell fate through regulation of a transcription factor (Yumerefendi et al. 2015) and the light induced nuclear exporter (LINX) to examine the rapid kinetics of deposition and removal of H2Bub1, which controls the trans-histone regulated methylation events of H3K4 and H3K79 (Yumerefendi et al. 2016). In this study, we applied LANS to precisely control Set2 localization, and consequently its activity, in order to quantitatively evaluate the dynamics of H3K36 methylation and demethylation.

We find that LANS-Set2 nuclear import results in rapid deposition of H3K36me2/3 (*t*_1/2_ = 20 min for H3K36me2 and *t*_1/2_ = 27 min for H3K36me3). Interestingly, although final H3K36me2/3 levels correlate with increased RNA abundance, as expected, the relative H3K36me3 rate of deposition over time is consistent between genes, regardless of transcriptional frequency. H3K36 demethylation upon Set2 nuclear export is also rapid (*t*_1/2_ = 40 min for H3K36me2 and *t*_1/2_ = 49 min for H3K36me3) and is largely regulated by Rph1. Intriguingly, the relative rate of H3K36me3 loss is largely uniform across all transcribed genes and mostly independent of RNA abundance, suggesting that H3K36me removal is largely uncoupled from transcription elongation. Together, these findings demonstrate the potential for optogenetic tools, coupled with high-throughput genomics approaches, to uncover key insights into the regulatory dynamics of histone PTMs.

## RESULTS

### Optogenetic control of Set2 cellular localization

To quantitatively explore methylation dynamics, we sought to generate a photoresponsive variant of Set2 (LANS-Set2) capable of reversible translocation into and out of the nucleus in response to blue light (Figure 1A). We inactivated a putative bipartite nuclear localization signal (NLS) in Set2 (residues 538-539 and 549-551) by mutating the lysines and arginines in the motif to glycines and serines to generate Set2_NLSΔ_ (Kosugi et al. 2009). We reasoned that these mutations, distal from the functional SET and SRI (Set2-Rpb1 interacting) domains (Figure S1A), would impact Set2 localization without affecting its catalytic activity. We then tested whether Set2 could be constitutively inactivated by fusing it to a nuclear export signal (NES) sequences (Figure S1B) (Yumerefendi et al. 2015). Expressing these proteins in a *SET2* deletion strain (set2Δ; Figure S1C) resulted in reduced H3K36 trimethylation and varying H3K36 dimethylation depending on the NES. Importantly, fusion of Set2_NLSΔ_ with the NLS in LANS (NLS-Set2_NLSΔ_) restored H3K36 methylation, although to less than wild-type levels (Figure S1C).

**Figure 1:**
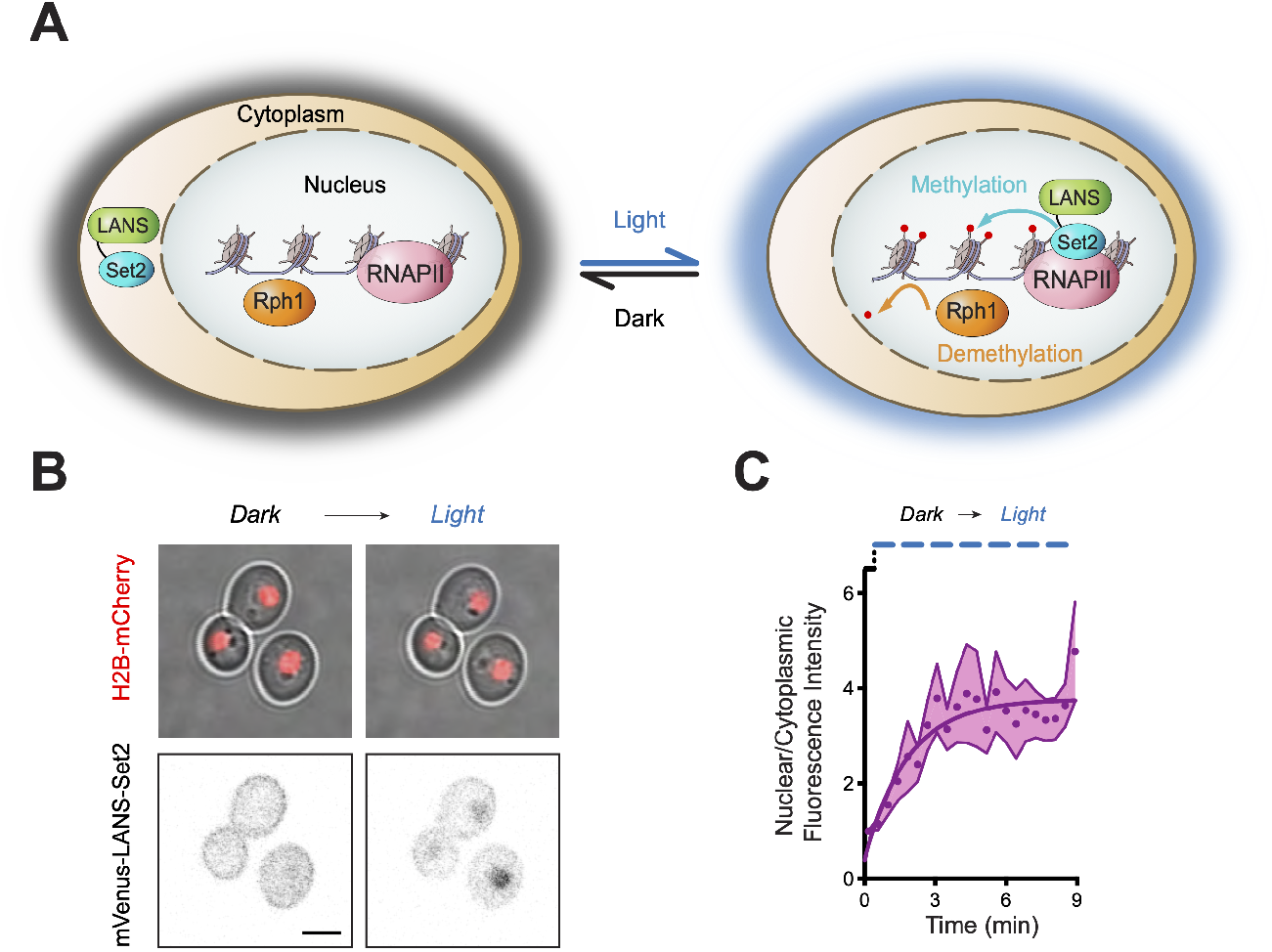
Optogenetic control of Set2 cellular localization. (A) Schematic of histone H3 lysine 36 (H3K36) methylation triggered by light-induced translocation of LANS-Set2 into the nucleus as well as demethylation by Rph1. (B) Confocal images from Additional file 1 demonstrating reversible control of mVenus-tagged LANS-Set2 localization in yeast cells with histone H2B endogenously tagged with mCherry (scale bar, 3 *μ*m). (C) Quantification of nuclear/cytoplasmic fluorescence intensity change before and during light activation. Mean ± SEM was calculated from the activation of multiple cells (*n* = 3) shown in (B) and Additional file 1.

We then expressed fluorescently tagged Set2, Set2_NLSΔ_, NES1-Set2_NLSΔ_, or NLS-Set2_NLSΔ_ in H2B-mCherry expressing cells to visualize subcellular localization of these static, non-shuttling constructs (Figure S1D). In contrast to wild-type Set2, which is nuclear, NLS-inactivated Set2 localized to both the nucleus and the cytoplasm. NES1-Set2_NLSΔ_ localized to the cytoplasm, and NLS-Set2_NLSΔ_ restored nuclear localization. These results further supported successful identification and elimination of the Set2 NLS and identified both an NES and NLS suitable for optogenetic control. Next, we combined these elements to generate a functional photoswitch. We expressed the NLS-mutated Set2 fused to a LANS variant using NES1 (mVenus-LANS-Set2) in the H2B-mCherry yeast strain and monitored nucleocytoplasmic ratios upon blue light exposure using confocal microscopy. Light exposure produced, on average, a three-and-a-half-fold change in nucleocytoplasmic ratio (Figures 1B-C, Additional file 1) with most Set2 entering the nucleus in less than 5 minutes. Thus, we identified NES1 as suitable for use in LANS-Set2 to achieve optimal light-induced nuclear translocation.

### LANS-Set2 regulates H3K36 methylation levels and Set2-associated phenotypes

Following removal of mVenus, which resulted in more stable Set2 protein levels (Figure S1E), LANS-Set2 was expressed in *set2*Δ cells. H3K36 methylation was evaluated following growth in the dark or light (Figure 2A). H3K36me3 and H3K36me2 were increased 6- and 4-fold, respectively, from the dark-to-light, whereas H3K36me1 levels were unchanged (Figures 2B and S1E).

**Figure 2:**
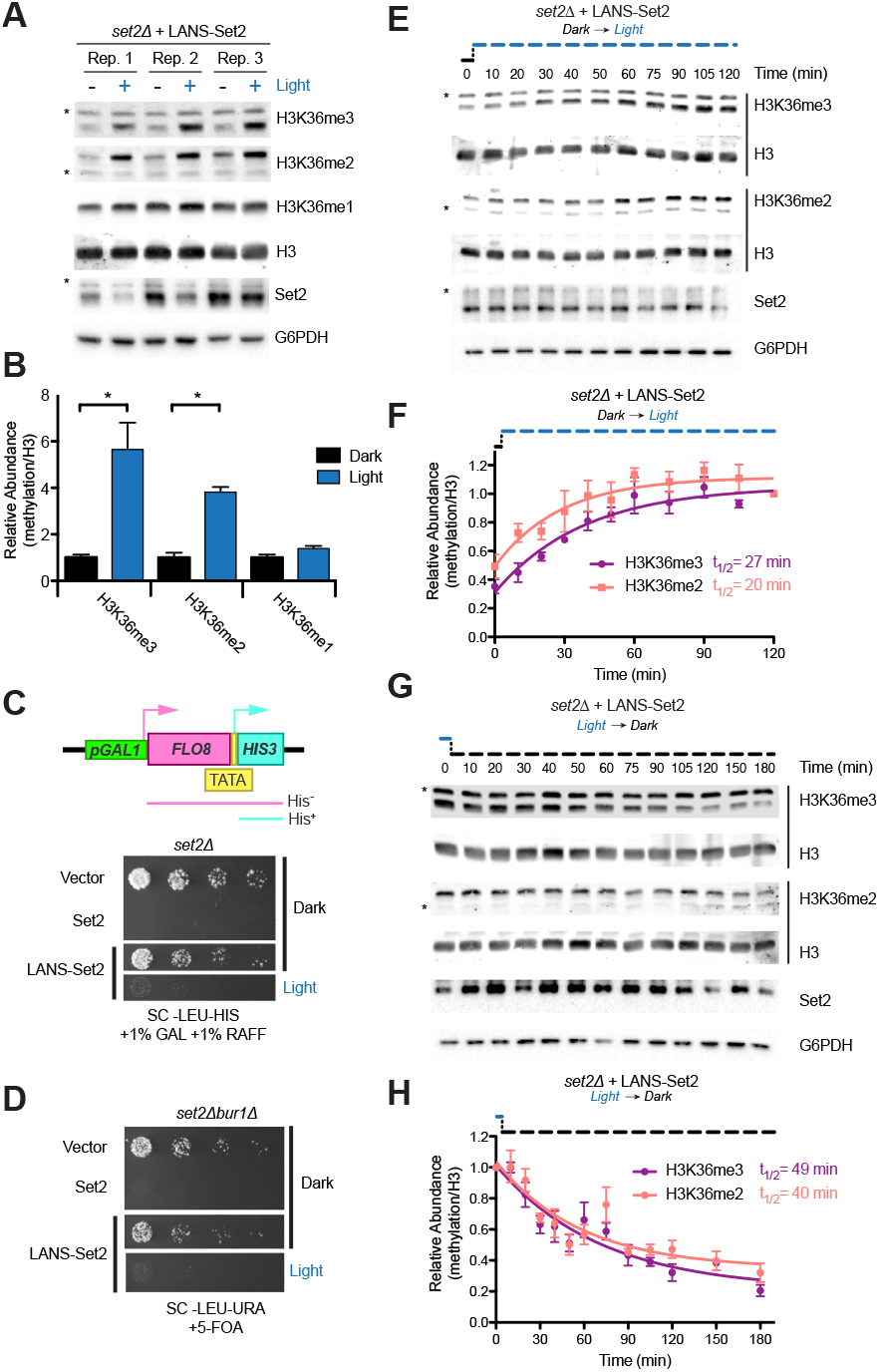
LANS-Set2 regulates H3K36 methylation levels and Set2-associated phenotypes. (A) Western blot analysis comparing levels of H3K36 methylation in whole cell lysates prepared from log phase cultures grown continuously in the dark or light. Asterisks indicate nonspecific bands. (B) Quantification of histone modifications from immunoblots in (A). Data represent mean values ± SD (*n* = 3). (C) Diagram of the *FLO8-HIS3* reporter. The promoter upstream of the *FLO8* gene has been replaced by a galactose inducible promoter and a *HIS3* cassette has been inserted out of frame from the *FLO8*_+1_ ATG such that growth in the absence of histidine can only occur when transcription initiates from an internal TATA located at *FLO8*_+1626_. (D) Four-fold serial dilutions of overnight *set2*Δ cultures expressing one of several constructs were spotted on the indicated solid media, which were incubated in the dark or light for 4 days before imaging (see Figure S1F for original images). LANS-Set2 phenocopies *set2*Δ in the dark and wild-type Set2 in the light. (E) Five-fold serial dilutions of overnight cultures of wild type BY4742 and *BUR1* plasmid shuffling strains were spotted on the indicated solid media, which were incubated in the dark or light for 3 days before imaging (see Figure S1G for original images). LANS-Set2 phenocopies *set2*Δ*bur1*Δ in the dark and wild-type Set2 in the light. (F) Representative western blot analysis of whole-cell lysates probing gain of H3K36 methylation over time using LANS-Set2 in *set2*Δ after the transition of log phase cultures from dark to light (see Figure S2A for replicates). Asterisks indicate nonspecific bands. (G) Quantification of H3K36 modifications as a function of time from triplicate immunoblots shown in (F) and Figure S2A. n = 3 and data represent mean ± SEM. (H) Representative western blot analysis of whole-cell lysates probing loss of H3K36 methylation over time using LANS-Set2 in *set2*Δ after the transition of log phase cultures from light to dark (see Figure S2B for replicates). Asterisks indicate nonspecific bands. (I) Quantification of H3K36 modifications as a function of time from triplicate immunoblots shown in (H) and Figure S2B. *n* = 3 and data represent mean ± SEM. Half-lives were calculated from single exponential fits to the H3K36me3 and H3K36me2 relative abundance data using GraphPad Prism 5. \**P* < 0.05.

We then evaluated the functional effects of LANS-Set2. LANS-Set2 was introduced into the KLY78 *set2*Δ strain in which *H1S3* was placed under the control of a cryptic promoter at *FLO8*_+1626_. Survival of these cells on solid media lacking histidine is dependent on conditions that enable aberrant transcriptional initiation from the cryptic promoter (Figure 2C) (Silva et al. 2012). Set2 loss promotes cryptic initiation at the *FLO8* locus. Colony growth assays showed that cells expressing LANS-Set2 grown in the dark phenocopy a *set2*Δ strain, whereas the same cells grown in the light phenocopy wild-type SET2, with some residual growth (Figures 2C and S1F). We also characterized LANS-Set2 in the *set2*Δ*burl*Δ *BUR1* shuffle strain wherein growth on solid media with 5-fluoroorotic acid (5-FOA), which selects against the *BUR1/URA3* plasmid, indicates the bypass of the requirement for Bur1 kinase (Keogh et al. 2005, 2003). The *bur1*Δ bypass spotting assay showed that cells expressing LANS-Set2 grown in the dark phenocopy a *set2*Δ strain, whereas these cells grown in the light phenocopy wild-type SET2 (Figures 2D and S1G). Taken together, these data indicate that LANS-Set2 modulates phenotypic effects associated with Set2 status.

We next interrogated the dynamics of H3K36 methylation and demethylation by LANS-Set2 in *set2*Δ cells. Cells were cultured in the dark or light until log phase growth. Light conditions were then reversed, and cells were collected over time to quantify H3K36me2 and H3K36me3. Following dark-to-light transition, in which LANS-Set2 localizes to the nucleus, we observed rapid accumulation of H3K36me2 (*t*_1/2_ = 20 min) and H3K36me3 (*t*_1/2_ = 27 min) (Figures 2E-F and S2A). Following light-to-dark transition, leading to export of LANS-Set2 from the nucleus, we observed loss of H3K36me2 (*t*_1/2_ = 40 min) and H3K36me3 (*t*_1/2_ = 49 min) (Figures 2G-H and S2B). To validate these kinetics, we employed a different method for nuclear depletion (Haruki et al. 2008). Anchor-away (AA) fuses a C-terminal FRB domain to Set2 at its native locus. Exposure to rapamycin results in the nuclear export of Set2-FRB. We probed for steady-state H3K36me levels with and without rapamycin. We also evaluated methylation over time following addition of rapamycin (Figures S2C-F). Although differences in steady-state H3K36me2 and H3K36me3 levels were greater for the AA method than for LANS-Set2 (Figures 2B and S2F), kinetics for loss of each mark were similar or slower for the AA method (Figures S2G-H). Despite overexpression of LANS-Set2 compared to Set2-FRB (Figures S2I-J), our optogenetic switch exhibits rapid kinetics and reversibility (Additional file 1), though with a lower dynamic range than the AA method.

Thus, our photoactivatable LANS-Set2 provides a rapid and reversible tool with which to probe dynamics of H3K36me gain and loss. Herein, LANS-Set2 activation refers to nuclear localization of LANS-Set2 (dark-to-light transition), and LANS-Set2 inactivation refers to cytoplasmic localization of LANS-Set2 (light-to-dark transition).

### Genome-wide examination of H3K36me3 reveals rapid methylation and demethylation kinetics

To interrogate the genome-wide dynamics of H3K36me3 gain and loss, we performed chromatin immuno-precipitation for H3K36me3 and H3 followed by high-throughput sequencing (ChIP-seq) in *set2*Δ cells expressing LANS-Set2. Cells were collected at multiple time points following either LANS-Set2 activation (0, 20, 40, 60 minutes), or LANS-Set2 inactivation (0, 30, 60, 90 minutes), as determined from the methylation kinetics detected by immunoblot (Figures 2F and 2H), constituting a longitudinal study design with replicate observations (*n* = 3). To enable quantitative normalization for subsequent analyses, ChIP-seq experiments were spiked-in with *Schizosaccharomyces pombe* (*S. pombe*) chromatin. H3K36me3 signal was normalized to H3 signal and scaled by the S. *pombe* reads. Overlapping genes were excluded from analyses as ChIP-seq signal could not be confidently attributed to an individual gene. To assess whether introns had significantly different H3K36 methylation patterns that could bias downstream analyses, we compared the mean signal of each intron to its flanking exons. Interestingly, though median intronic signal was lower than median exonic signal, this difference did not reach significance at any timepoint across all replicates in LANS-Set2 activation (Figures S3A-D). Fold change between intronic and pre-exonic signal also lacked a clear trend, appearing normally distributed with a mean around 0 (Figures S3E-H). This pattern was consistent in the context of LANS-Set2 inactivation (Figures figS3I-P). Based on these results, intron-containing genes were not excluded from subsequent analyses.

As expected, H3K36me3 signal primarily localized over gene bodies, rather than the transcription start sites (TSSs) of genes (Figure 3A). There was also a clear increase and decrease in H3K36me3 levels following LANS-Set2 activation and inactivation, respectfully (Figure 3A). To determine how H3K36me3 temporal dynamics varied across the genome, we first calculated the average H3K36me3 signal at each timepoint for every gene across each of the replicates (*n* = 3) following LANS-Set2 activation and inactivation. Consistent with our western blots, H3K36me3 signal increased and decreased at genes following LANS-Set2 activation and inactivation, respectfully (Figures 3B-C). H3K36me3 changes occurred relatively equally over genes without a 5’ or 3’ preference (Figures S3Q-R). Per-gene H3K36me3 signal increased at an approximately linear rate following LANS-Set2 activation (Figure 3B). In contrast, H3K36me3 signal loss was less linear; individual gene H3K36me3 signal loss following LANS-Set2 inactivation occurred largely within the first 60 minutes.

**Figure 3:**
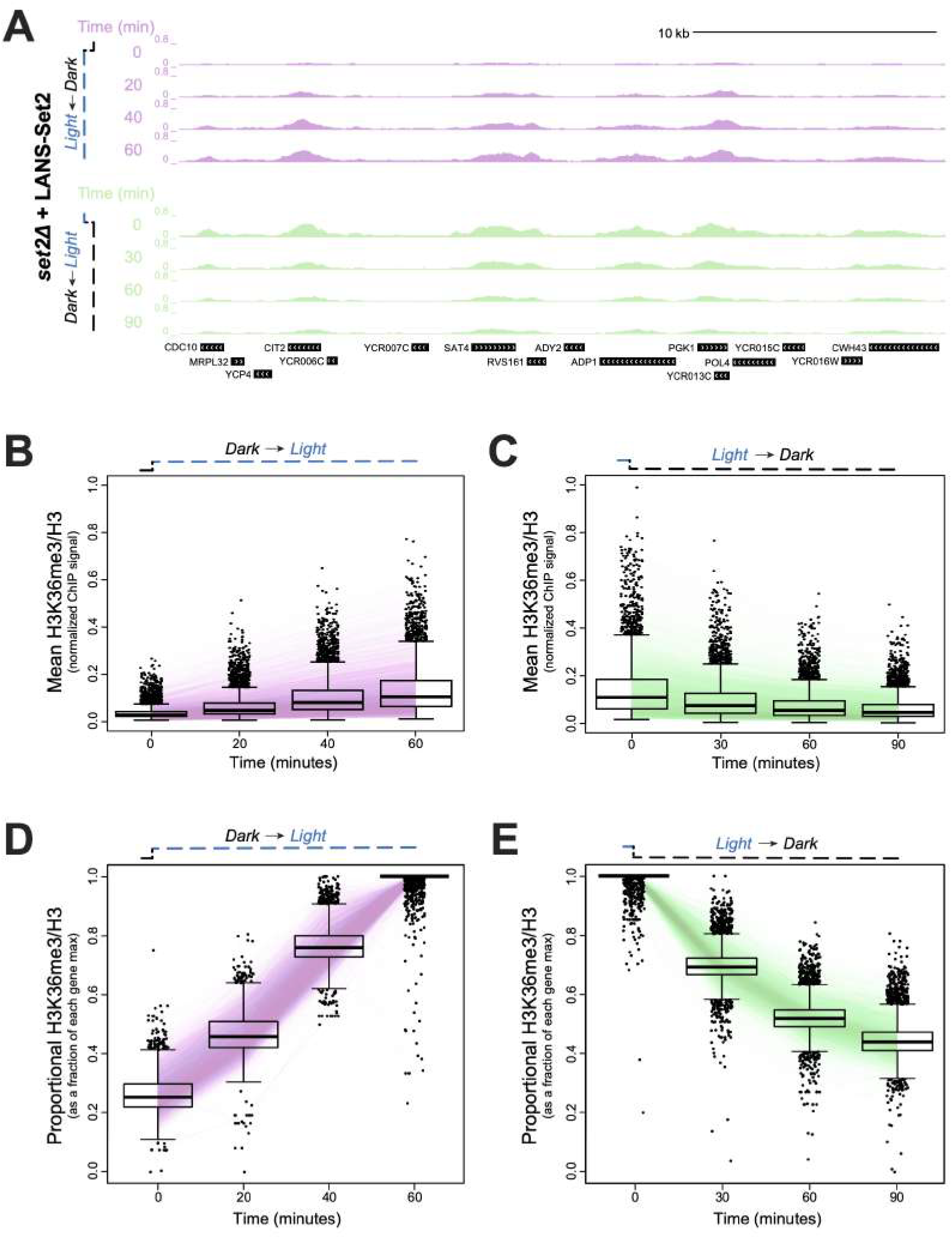
Genome-wide examination of H3K36me3 reveals rapid methylation and demethylation kinetics. (A) Genome browser ChIP-seq signal track of a representative example of LANS-Set2 activation (green) and inactivation (purple) over the time course experiment. Signal is normalized by H3 ChIP-seq signal and scaled by the internal spike-in *S. pombe* DNA. (B,C) Distribution of mean, per-gene normalized LANS-Set2 activation (B) or LANS-Set2 inactivation (C) ChIP-seq signal represented as interquartile range boxplots over the time course. Each line represents the mean of the replicates for a specific gene over time. (D,E) Distribution of the relative (D) LANS-Set2 activation or (E) LANS-Set2 inactivation ChIP-seq signal over time. To highlight relative H3K36me3 changes for each gene, the maximum signal value over all timepoints was set to 1, and subsequent time point became a fraction of that maximum.

Previous studies (Pokholok et al. 2005) identified an association between H3K36me3 levels at genes and RNA abundance, which may explain the variations seen in mean H3K36me3 ChIP-seq levels. The broad differences in H3K36me3 levels between genes obscured our ability to discern changes in the subtle patterns and/or trends of H3K36me3 deposition or removal over time (Figure 3C). To account for this, we scaled the H3K36me3 signal for each gene relative to the gene maximum H3K36me3 signal per replicate, resulting in H3K36me3 signal represented as a fraction of maximum H3K36me3 signal. By doing this, we created an internally normalized, per-gene H3K36me3 signal for each timepoint (henceforth referred to as relative H3K36me3 signal). Normalizing each gene relative to itself allowed us to directly compare the rates of H3K36me3 deposition or removal between genes independent of overall H3K36me3 levels.

As expected, the maximum H3K36me3 signal for most genes was observed at the final timepoint (for LANS-Set2 activation) and initial timepoint (for LANS-Set2 inactivation; Figure 3D-E, represented as a fractional value of 1). This approach highlighted the linearity of H3K36me3 signal gain upon LANS-Set2 activation. In contrast, H3K36me3 loss was non-linear with a rapid decrease over the initial 0-60 minutes, slowing over the subsequent 30 minutes. Although the final H3K36me3 levels for a specific gene reflects transcript abundance, these data suggest that the relative gain of methylation occurs at a consistent rate for most genes. In contrast, loss of methylation resembles an exponential decay trend. Considering H3K36 methylation state as the product of enzymatic activities suggests that H3K36me3 deposition is rate limited by an external factor whereas H3K36me3 removal occurs by a more stochastic mechanism.

### A Bayesian generalized linear mixed effect model for H3K36me3 dynamics defines fixed and stochastic properties of H3K36me3 gain and loss

We next sought to understand whether H3K36me3 dynamics could be attributed to underlying biological differences observed by applying a statistical model that also accounted for intra-gene variability in the relative H3K36me3 gain and loss over time. We modeled the temporal dynamics of H3K36me3 signal for each gene, seeking to detect patterns consistent across all the genes considering all replicates. We used a Bayesian generalized linear mixed effect model (GLMM), which simultaneously accommodates the non-normality of the H3K36me3 signal (as either quasi-counts or quantiles) and leverages the longitudinal study design, nature of the data, and replicates to identify genes with significant gain or loss of H3K36me3 signal with time (Figures 4A and S4A-B).

**Figure 4:**
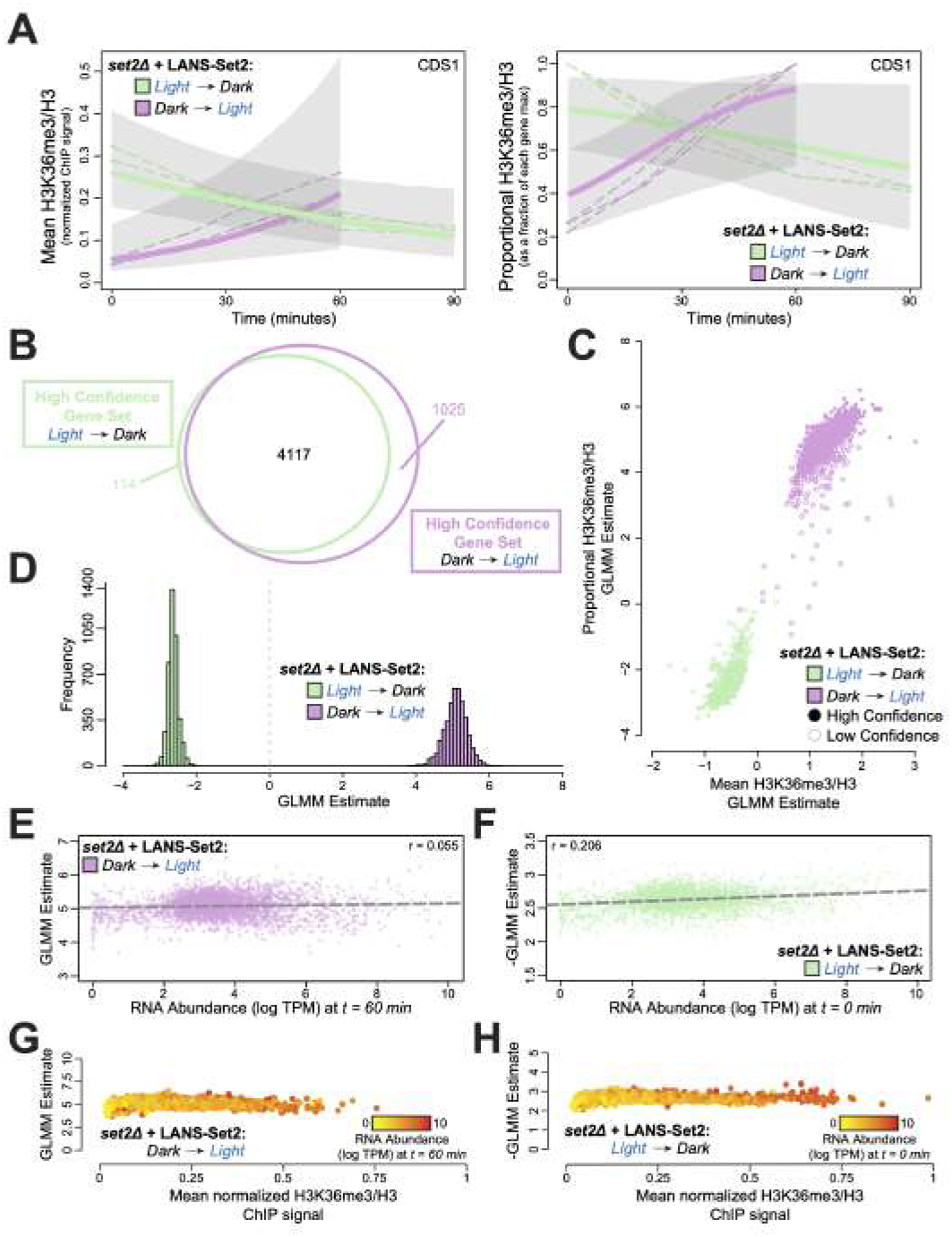
A Bayesian generalized linear mixed effect model for H3K36me3 dynamics defines fixed and stochastic properties of H3K36me3 gain and loss. (A) Posterior H3K36me3 rates from Bayesian generalized linear mixed model (GLMM) for normalized ChIP-seq signal (left) and relative H3K36me3 ChIP-seq signal (right) for the gene CDS1 (YBR029C) throughout the timecourses of LANS-Set2 activation (green) and LANS-Set2 inactivation (purple). Dashed lines represent individual ChIP-seq replicates, while bold lines represent the GLMM posterior mean of the rate. Shaded regions indicate the 95% credible interval on the rate parameter. (B) Venn diagram of the high confidence genes identified within LANS-Set2 activation (green) and LANS-Set2 inactivation (purple). High confidence genes had a clear positive or negative rate, defined as having 95% credible intervals that never include zero (on the linear predictor). (C) Per-gene GLMM rates for normalized H3K36me3 ChIP-seq signal and relative H3K36me3 ChIP-seq signal for both LANS-Set2 activation (green) and LANS-Set2 inactivation (purple). Solid circles signify high confidence genes, while hollow circles represent low confidence genes. (D) Histogram of GLMM rates within the shared high confidence gene set between LANS-Set2 activation (green) and LANS-Set2 inactivation (purple) (*n* = 4117). (E) LANS-Set2 activation GLMM rates compared to mean RNA abundance levels (log TPM) at *t* = 60 minutes for genes that were high confidence for both LANS-Set2 activation and inactivation. The Pearson correlation coefficient is *r* = 0.055. The dashed lines represent the line of best fit, (F) LANS-Set2 inactivation GLMM rates compared to mean RNA abundance levels (log TPM) at *t* = 0 minutes for genes that were high confidence gene in both sets. Pearson correlation coefficient is *r* = 0.206. Dashed line represents the line of best fit. (G) LANS-Set2 activation GLMM rates compared to average, normalized H3K36me3 ChIP-seq levels at *t* = 60 minutes for each gene in the shared, high confidence gene set. Each gene is colored by mean RNA abundance levels (log TPM) at the same time point. (H) LANS-Set2 inactivation GLMM rates compared to average, normalized H3K36me3 ChIP-seq levels at *t* = 0 minutes for each gene in the shared, high confidence gene set. Each gene is colored by mean RNA abundance levels (log TPM) at the same time point.

Using the model, we defined a class of high confidence genes with significant temporal dynamics based on having a 95% highest posterior interval on the rate parameter that did not include 0. Using relative H3K36me3 signal, we found 4231 (79%) and 5142 (96%) high confidence genes (out of 5355 total genes) in the LANS-Set2 activation and inactivation, respectively. Of these, 4117 were high confidence in both LANS-Set2 activation and inactivation (Figures 4B and S4C). Modeling the H3K36me3 data generated consistent trends using either absolute or relative H3K36me3 signal (quasi-counts; see Methods), suggesting that our modeling approaches were not biased based on either data transformation (Figures 4C and S4D). Relative LANS-Set2 activation rates were more extreme than inactivation rates suggesting, across genes, H3K36me3 is deposited faster rate than it is lost (Figures 4D and S4E). This result is consistent with our western blots (Figures 2F and 2H). Relative H3K36me3 gain rates correlated with loss rates (*r* = 0.438) implying, in general, genes that are rapidly methylated are also rapidly demethylated (Figure S4F).

We asked whether genomic features could account for variability in the rate of H3K36me3 gain and loss across genes. We evaluated gene length, average H3K36me3 levels, and RNA abundance levels. RNA-seq (*n* = 3) was performed at each ChIP-seq time point in both LANS-Set2 activation and inactivation conditions (however, one replicate failed at the 60-minute timepoint in LANS-Set2 activation). Consistent with previous studies (Kemmeren et al. 2014; McDaniel and Strahl 2017), the loss of Set2 had a relatively limited impact on RNA abundance (Figure S4G). The RNA abundance for 445 genes (out of 6692) was significantly different after LANS-Set2 activation (Figure S4H). LANS-Set2 inactivation affected the RNA of 313 genes (Figure S4I). Interestingly, LANS-Set2 activation was associated predominantly with increased RNA abundance (313 genes with increased levels vs 132 decreased), whereas LANS-Set2 inactivation decreased more (245 genes decreased vs 68 increased). Following LANS-Set2 activation, enriched RNA ontologies included “oxidoreductase activity” (adj. *p* ≤ 4.96 × 10^−11^) for genes with decreased RNA abundance, and “structural constituent of ribosome” (adj. *p* ≤ 2.40 × 10^−73^) for genes with increased abundance. Genes with increased RNA abundance after LANS-Set2 inactivation were not significantly enriched for any specific ontologies, however genes that decreased had one significant ontology, “oxidoreductase activity” (adj. p ≤ 1.78×10^−6^).

Before associating relative H3K36me3 rates of change with genomic features, we explored the relationship between H3K36me3 levels and RNA abundance. Maximum LANS-Set2 levels (*t* = 60 min in LANS-Set2 activation, *t* = 0 min in LANS-Set2 inactivation) were highly concordant with RNA abundance (*r* = 0.467 and *r* = 0.486 respectively, Figure S4J) in the high confidence gene set, confirming prior studies (McDaniel and Strahl 2017). We also compared gene length to both mean H3K36me3 signal and RNA abundance at timepoints most closely resembling wild-type conditions, and found limited (Figure S4K) or no (Figure S4L) association for both Set2 activation and inactivation conditions.

We then asked whether gene length or H3K36me3 levels associates with the relative rate of H3K36me3 change. Surprisingly, neither feature was predictive of relative H3K36me3 rates of change after LANS-Set2 activation (Figures 4E and S4M-P) or inactivation (Figures 4F and S4Q-T). We therefore hypothesized that H3K36me3 rates of change may be regulated through transcriptional processes, and specifically investigated whether H3K36me3 change was associated with RNA abundance. Surprisingly, RNA abundance was not strongly correlated with the relative rate of H3K36me3 gain upon Set2 activation (Figure 4G, *r* = 0.055), though H3K36me3 loss rates with LANS-Set2 inactivation were slightly correlated (Figure 4H, *r* = 0.206).

### H3K36me3/2 removal is rapid and largely mediated by Rph1 and Jhd1

We next investigated to the relative impact of putative demethylases on H3K36 dynamics. In a LANS-Set2 expressing *set2*Δ strain, we deleted putative demethylases in yeast (Rph1, Jhd1, Ecm5, Gis1) (Tu et al. 2007). Cells were grown in the light and probed for H3K36me3 and H3K36me2. *RPH1* deletion had the largest effect on H3K36me3, increasing global levels by ~2-fold (Figures 5A-B, S5A-B). Without using LANS-Set2, deletion of *RPH1* in the context of wild-type Set2 increases H3K36me3 by ~1.3-fold (Figures S5C-D).

**Figure 5:**
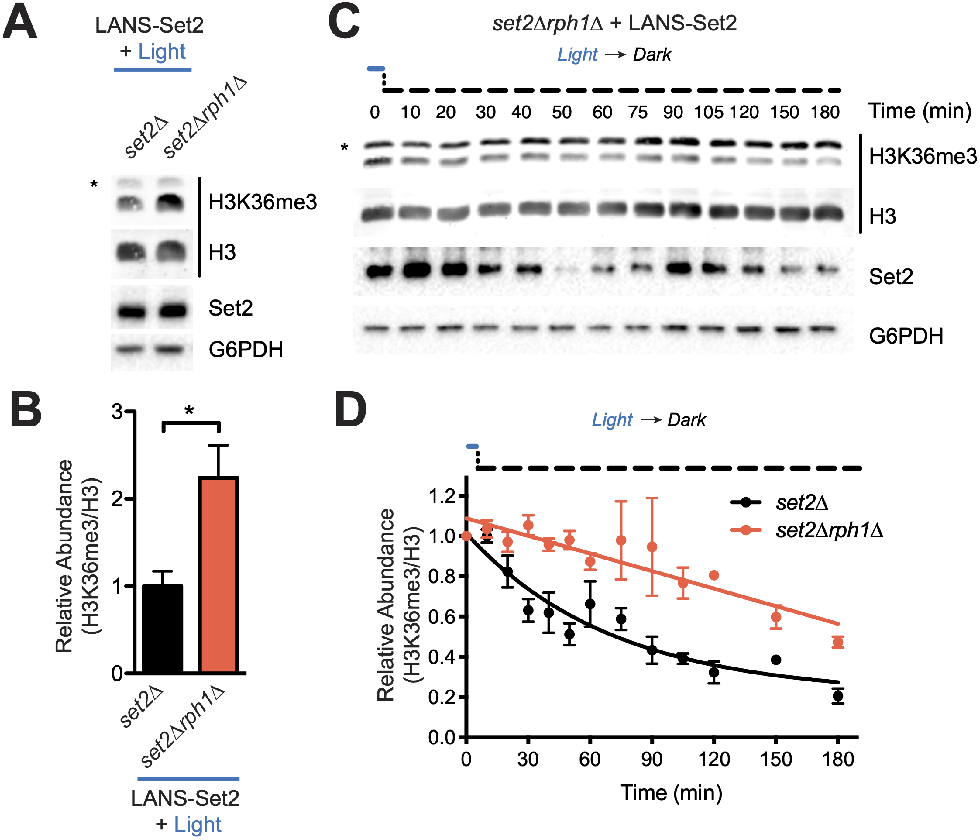
H3K36me3/2 removal is rapid and largely mediated by Rph1 and Jhd1. (A) Representative western blot analysis of whole cell lysates prepared from log phase cultures of the indicated strains transformed with LANS-Set2 and grown continuously in the light. Asterisks indicate nonspecific bands. (B) Quantification of H3K36me3 from triplicate immunoblots shown in (A) and Figure S6A. Data represent mean values ± SD (*n* = 3). (C) Representative western blot analysis of whole-cell lysates probing loss of H3K36me3 over time using LANS-Set2 in *set2*Δ*rph1*Δ after the transition of log phase cultures from light to dark (see Figure S6E for replicates). Asterisks indicate nonspecific bands. (D) Quantification of H3K36me3 as a function of time from triplicate immunoblots shown in Figures 2H and S2B (*set2*Δ) and Figures 5C and S5E (*set2*Δ*rph1*Δ). *n* = 3 and data represent mean ± SEM. The half-life was calculated from a single exponential fit to the H3K36me3 relative abundance data using GraphPad Prism 5. \**P* < 0.05.

The dynamics of H3K36 demethylation were then evaluated following LANS-Set2 inactivation. We found that the rate of H3K36me3 loss was most impacted in the *set2*Δ*rph1*Δ strain (compare Figure 2H to Figures S5F, S5H, S5J, and S5L). The rate of loss of H3K36me2 was most impacted in the *set2*Δ*jhd1*Δ and *set2*Δ*ecm5*Δ strains, and the rate of loss in the *set2*Δ*gis1*Δ strain was similar to the *set2*Δ strain (Figures S5G-L). We also examined whether disruption of *ASF1*, a histone exchange factor that contributes to replication and transcription, would affect loss of methylation. Deletion of *ASF1* minimally impacted loss of H3K36 methylation (Figures S5M-P) (Rufiange et al. 2007). We also examined the impact of replication by adding α-factor to arrest cells in G1 prior to LANS-Set2 inactivation. Cell cycle arrest minimally impacted loss of H3K36 methylation (Figures S5Q-T). Taken together, these data indicate that Rph1 primarily mediates loss of H3K36 trimethylation whereas Jhd1 is primarily responsible for H3K36me2 demethylation.

### Global H3K36me3 removal is associated with a one phase exponential rate of decay

To explore the impact of deregulated demethylation on H3K36me3 distribution, we performed ChIP-seq time course following LANS-Set2 inactivation in *set2*Δ*rph1*Δ cells. In the absence of Rph1, H3K36me3 signal was retained over genes, and total H3K36me3 levels were increased (Figures 6A and S6A). Th effect of Rph1 loss was apparent when comparing all genes across the timepoints (Wilcoxon rank sum test, *p* < 2.2 × 10^−16^ for all time points, Figure 6B). Following scaling of the average H3K36me3 levels relative to each gene’s maximum H3K36me3 level, we observed a clear difference between *set2*Δ*rph1*Δ and *set2*Δ relative H3K36me3 signals, with the largest difference between strains at 60 minutes following LANS-Set2 inactivation (Figure 6C). That signal differences between the conditions were largely eliminated by 90 minutes suggests that Rph1 is most active in the early demethylation of H3K36me3.

**Figure 6:**
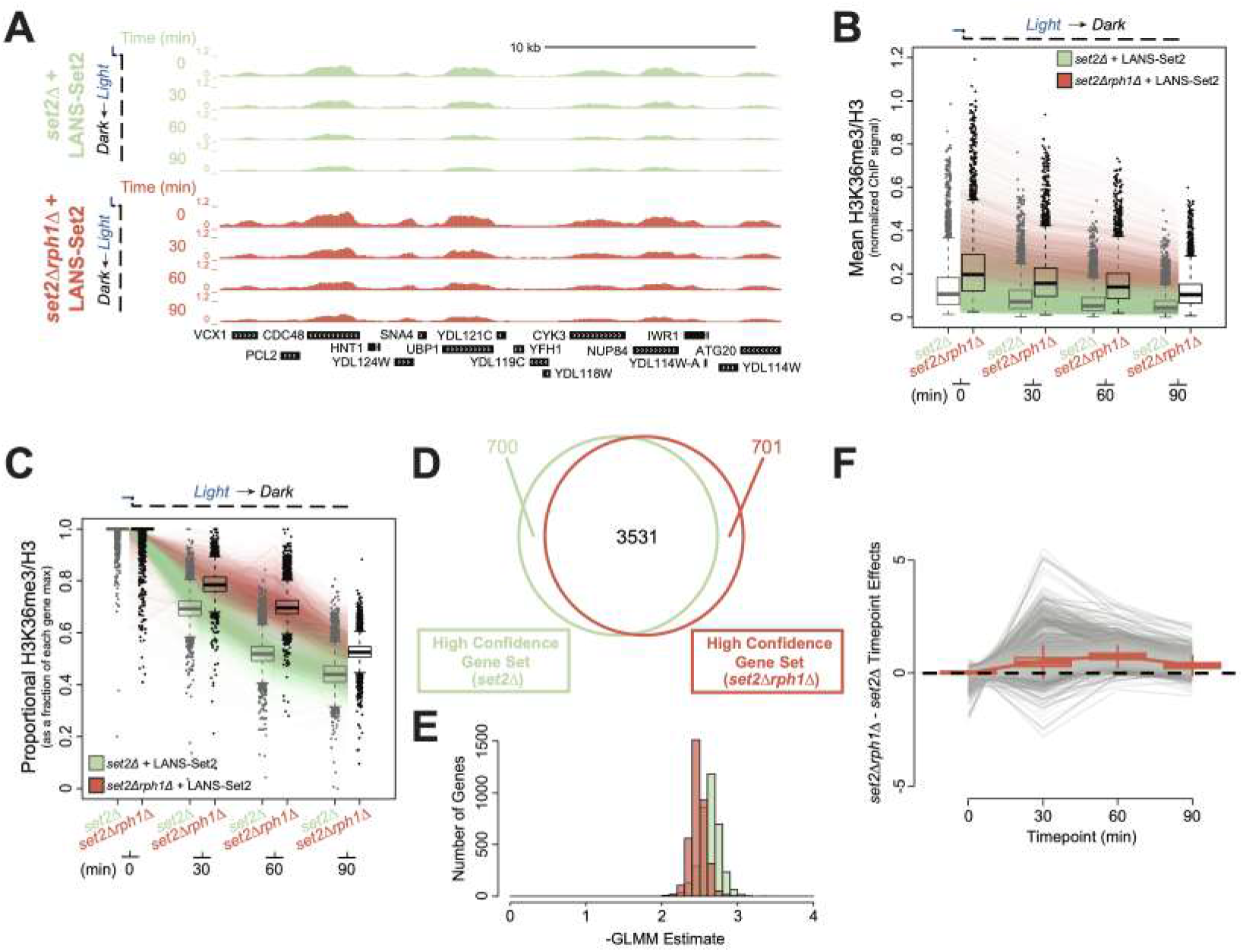
Global H3K36me3 removal is associated with a one phase exponential rate of decay. (A) Genome browser ChIP-seq signal track of a representative example of LANS-Set2 inactivation in a *set2*Δ (green) or *set2*Δ*rph1*Δ (red) background the time course experiment. Signal is normalized by internal spike-in *S. pombe* DNA and H3 ChIP-seq signal. (B) Distribution of mean, per-gene normalized LANS-Set2 inactivation ChIP-seq signal represented as interquartile boxplots across the time course in a *set2*Δ (green) or *set2*Δ*rph1*Δ (red) background. Each line represents the mean of all three replicates for one specific gene over time. Gray boxplots represent LANS-Set2 inactivation in the *set2*Δ background (as seen in Figure 3C) while black boxplots represent the *set2*Δ*rph1*Δ background. (C) Distribution of the relative LANS-Set2 inactivation ChIP-seq signal in a *set2*Δ (green) or *set2*Δ*rph1*Δ (red) over time. To highlight relative H3K36me3 change for each gene, the maximum signal value over all timepoints was set to 1, and subsequent time point became a fraction of that maximum. Gray boxplots represent LANS-Set2 inactivation in the *set2*Δ background (as seen in Figure 3E) while black boxplots represent the *set2*Δ*rph1*Δ background. (D) Venn diagram of the high confidence genes identified within LANS-Set2 inactivation in a *set2*Δ (green) or *set2*Δ*rph1*Δ (red) background. High confidence genes were determined to have a clear negative trend based on the 95% credible interval of the GLMM rate never including zero (on the linear predictor). (E) Histogram of the GLMM rates within the shared high confidence gene set between LANS-Set2 inactivation in a *set2*Δ (green) or *set2*Δ*rph1*Δ (red) background (*n* = 3531). (F) Jointly modeling the *set2*Δ and *set2*Δ*rph1*Δ backgrounds of LANS-Set2 inactivation rates revealed a lag in *set2*Δ*rph1*Δ compared to *set2*Δ across the time course. Points above zero indicate genes with a higher GLMM loss rates in *set2*Δ compared to *set2*Δ*rph1*Δ at a given timepoint, while points below zero indicates genes with lower GLMM rates in *set2*Δ*rph1*Δ compared to set2Δ. Each line represents one gene across the time course. Boxplot borders represent the interquartile range of the difference in GLMM rates between *set2*Δ*rph1*Δ and *set2*Δ for each timepoint.

We then applied GLMM to the *set2*Δ*rph1*Δ data (Figures S6B-C). Using the estimated relative H3K36me3 rates of loss, we identified 3531 (out of 5355 total) high confidence genes after Set2 inactivation across both *set2*Δ*rph1*Δ and *set2*Δ strains (Figure 6D). We observe that relative *set2*Δ*rph1*Δ H3K36me3 loss rate estimates were, as a whole, less extreme than the relative *set2*Δ H3K36me3 loss rates (Figure 6E), supporting that H3K36me3 is demethylated in *set2*Δ*rph1*Δ cells more slowly than *set2*Δ alone. Modeling the *set2*Δ*rph1*Δ H3K36me3 relative loss rates directly against those in *set2*Δ demonstrated that H3K36me3 relative loss rates were significantly delayed in *set2*Δ*rph1*Δ versus *set2*Δ samples (Figure 6E). H3K36me3 signal loss became more disparate between 0 and 60 minutes, before partially recovering at 90 minutes. Together, these data suggest that Rph1 loss significantly slows H3K36me3 demethylation by primarily mediating early demethylation prior to other factors.

Finally, we examined the relationship of H3K36me3 decay in the absence of Rph1 to RNA abundance. Relative H3K36me3 demethylation rates in the absence of Rph1 were not strongly correlated to transcriptional frequency (Figure S6D; *r* = 0.132), suggesting that H3K36me3 loss occurs uniformly at genes in a stochastic manner.

## DISCUSSION

In this study we sought to quantitatively explore the dynamics of H3K36me addition and removal in response to Set2 modulation. We created a light-controlled variant of Set2 (LANS-Set2) that offered a rapid and reversible tool. Using this photoswitch, we found that H3K36me3 deposition by Set2 is rapid. Total levels positively correlate with transcriptional frequency, in agreement with prior studies. Intriguingly, however, by internally scaling the rate of H3K36me3 deposition based the total H3K36me3, we found the rate at which each gene achieves its maximum level to be largely uniform. These data suggest that H3K36me3 deposition is regulated, perhaps through the activity of RNA polymerase, which may associate with a finite number of Set2 molecules during each round of transcription. In this model, a higher level of H3K36me3 at more highly transcribed genes results from multiple rounds of transcription, in which each round of transcription mediates a fixed amount of H3K36 methylation per nucleosome across the population of cells. Additionally, the linear rate of H3K36me3 gain across protein-coding genes also suggests that Set2 activity remains constant over the time course of H3K36 methylation deposition. H3K36me deposition may be controlled by a limited amount of Set2 that can interact with RNAPII. Alternatively, the residency time of Set2 at nucleosomes could be influenced by the speed of RNAPII elongation. Consistent with the idea of limited binding capacity of Set2 with RNAPII at genes, we observed that overexpression of *SET2* in yeast does not result in increased H3K36me levels (DiFiore *et al.*, in press at *Cell Reports*). Future studies will be required to explore this model further, for example, by extending either the length of the CTD or levels of serine 2 CTD phosphorylation as a means to accommodate additional molecules of Set2 on RNAPII.

In contrast to H3K36me addition, H3K36me2/3 demethylation occurs more slowly than deposition. Demethylation exhibits a pattern that cannot be explained by simple passive loss through cell division or by histone exchange. Rather, and in agreement with work by others, H3K36me2/3 is removed through the activity of multiple H3K36 demethylases, primarily Rph1 that targets H3K36me3. Intriguingly, we showed that H3K36me2/3 is lost at the same time scale for all genes, regardless of H3K36 methylation levels. This first order kinetic pattern of demethylation suggests that the rate of demethylation is regulated by the abundance and equal availability of H3K36 methylation as a substrate. This pattern differs from that associated with a mechanism of directed or controlled removal, which might be observed if a fixed amount H3K36 demethylases were associated with RNAPII during transcription elongation. In this scenario, highly transcribed genes would lose H3K36me3 more rapidly than lowly transcribed ones, and the rate at which H3K36me3 is lost would appear linear and correlate with RNA abundance levels. As relative H3K36me3 loss resembles a decay curve and we found minimal association between H3K36me3 loss rates and RNA abundance, our study suggests that H3K36 demethylation occurs through a stochastic and continuous mechanism.

Although Set2 methylation is known to function in transcription-associated activities, including the prevention of cryptic transcription, a recent report uncovered the potential for Set2 methylation and demethylation by Rph1 to function in metabolic pathways that control amino acid biosynthesis (Ye et al. 2017). More specially, the Tu lab showed that Set2 methylation promotes S-adenosylmethionine (SAM) consumption to drive cysteine and glutathione biosynthesis, whereas H3K36me removal is linked to the biosynthesis of methionine. Thus, H3K36 acts as a methylation “sink” that controls various biosynthetic pathways involving SAM. Consistent with this report, our studies support the idea that Rph1 acts in a global and stochastic manner for a role other than one in transcription. Our data, in combination with previous work showing that H3K36me2 is largely targeted by Jhd1 (Fang et al. 2007), may suggest that the combination of multiple H3K36 demethylases in fact function to maintain the balance of available SAM and the histone methylation sink.

The histone sink theory described above leads to an important question: how does Set2 methylation function in transcriptional regulation but also in amino acid biosynthetic pathways that are perhaps uncoupled to transcription? Although at first approximation it might seem these two functions of Set2 are incompatible, they may not be mutually exclusive. For example, although limited transcription is sufficient to regulate the Rpd3S deacetylation pathway, additional methylation offers the methyl sink. In this model, the high levels of H3K36me in highly transcribed genes would serve as a source for SAM regulation. Consistent with this idea, mutations in several transcription elongation factors like the PAF complex and Spt6, or Set2 itself, that limit H3K36 methylation are sufficient to prevent cryptic initiation (Cheung et al. 2008; Dronamraju and Strahl 2014; Silva et al. 2012). Furthermore, we observed that those genes that were the most impacted by Set2 loss were metabolic genes associated with amino acid biosynthesis.

In summary, our optogenetic system offers a level of control that permitted genome-wide analysis of H3K36me methylation and demethylation. By combining optogenetic, genomic and statistical techniques, we achieved fine resolution of dynamics of H3K36 methylation genome-wide. This provided inference on the parameters of transcription-dependent deposition and transcription-independent removal that builds from prior studies to illuminate new details on how the cycle of Set2 methylation and removal is achieved. These studies also offer a strategy for further use of optogenetic approaches to study other chromatin modifiers with improved spatiotemporal control, and to obtain a more quantitative, time-resolved understanding of the dynamics of chromatin regulation.

## METHODS

### Reagents

Antibodies: Set2 (raised in lab, 1:5000), G6PDH (Sigma Aldrich A9521, 1:100000), H3K36me3 (Abcam 9050, 1:1000 for ECL, 1:2000 for LI-COR and 2 *μ*L for ChIP), H3K36me2 (Active Motif 38255, 1:1000 for ECL and 1:2000 for LI-COR), H3K36me1 (Abcam 9048, 1:1000 for ECL and 1:2000 for LI-COR), H3K79me3 (Abcam 2621, 1:2000), H3 (Figure S1C EpiCypher 13-0001, 1:1000; Figure S1E Abcam 12079, 1:1000; CST 14269, 1:2000 for LI-COR and EMD Millipore 05-928 2 *μ*L for ChIP). Rabbit (Amersham NA934, Donkey anti-Rabbit), goat (Santa Cruz 2768, Rabbit anti-Goat), rabbit (Thermo SA5-10044, Donkey anti-Rabbit DyLight 800) and mouse (Thermo 35518, Goat anti-mouse DyLight 680) secondary antibodies were used at 1:10000.

### Strain generation

All strains were in the BY4741 background unless otherwise stated. Gene deletions (*SET2, RPH1, JHD1, ECM5*, and *GIS1*) were performed by gene replacement using the PCR toolkit. The Set2-FRB strain was generated by amplifying FRB-KanMX6 from pFA6a-FRB-KanMX6 (HHY168, Euroscarf) and inserting it at the *SET2* 3’ end by homologous recombination. Strains are listed in Supplementary Table 1.

### DNA Cloning

The mVenus-NES1-Set2 plasmid was Gibson assembled from an mVenus-NES1-MCS plasmid and *SET2* amplified from BY4741 genomic DNA. The resulting mVenus-NES1-Set2 plasmid was then blunt end cloned to create the mVenus-Set2 plasmid by digestion with XbaI and XmaI, polishing with Phusion polymerase and subsequent ligation to remove NES1. The mVenus-NES1-Set2 plasmid was also used to make the NES2 and NES3 variants by cutting the plasmid with XbaI and SbfI to remove NES1 and ligating annealed inserts.

Similarly, the mVenus-NES1-Set2_NLSΔ_ plasmid was Gibson assembled from the mVenus-NES1-MCS plasmid and *SET2*_*NLS*δ_ with XmaI restriction site at its 5’ and XhoI at its 3’ that was generated by two rounds of overlap extension PCR from BY4741 genomic DNA to sequentially mutate the bipartite NLS. The mVenus-NES1-Set2_NLSΔ_ plasmid was then used to make plasmids as above: the mVenus-Set2_NLSΔ_ plasmid was generated by blunt end cloning and the NES2, NES3 and NLS variants were generated by cutting and ligating annealed inserts.

The mVenus-LANS-Set2 plasmid was constructed by inserting *SET2*_*NLS*δ_ into the MCS of an mVenus-LANS-MCS plasmid: *SET2*_*NLS*δ_ was generated as above, both the insert and the plasmid were cut with XmaI and XhoI and ligation was performed. The resulting plasmid was used to generate the LANS-Set2 plasmid lacking mVenus: a LANS-Set2_NLSΔ_ cassette with HpaI restriction site at its 5’ and XhoI at its 3’ and the mVenus-LANS-Set2 plasmid were cut with HpaI and XhoI and ligated to remove mVenus. All plasmids were sequence verified (Eurofins). Selected plasmids are listed in Supplementary Table 2.

### Microscopy

Yeast samples were imaged and photo-activated with an Olympus FV1000 confocal microscope equipped with a 100 (N.A. 1.40) oil immersion objective. Image acquisition for mVenus and mCherry used the 514 nm and 559 nm laser lines, respectively, to acquire 800 × 800 images of a single optical section. For photo-activation of cells expressing mVenus-LANS-Set2, a timeline of image acquisition and photo-activation was generated with the Time Controller module in the Olympus Fluoview software. An image was taken before activation, after which activation images were taken every 25 seconds with activation in between each image acquisition for 10 cycles. After activation, images were acquired every 10 seconds for 50 cycles of imaging. The activation sequence consisted of rasterizing 800 × 800 pixels with 1% of the 488 nm laser and pixel dwell time of 8 *μ*s/pixel.

### Spotting

Overnight cultures of relevant strains were transformed and plated on appropriate SC plates that were then incubated at 30 °C in the dark. Colonies were resuspended in the appropriate SC dropout media and grown at 30 °C in the dark. Overnight cultures were diluted to an OD_600_ of 0.5 (*FLO8-HIS3* cryptic transcription initiation assay) or 2.0 (*bur1*Δ bypass assay). Six-fold (*FLO8-HIS3* assay) of five-fold (*bur1*Δ assay) serial dilutions were spotted onto appropriate plates. For the *GAL*-inducible *FLO8-HIS3* cryptic transcription initiation assay, dilutions were spotted onto SC–Leu–His plates containing 1% galactose and 1% raffinose as well as SC–Leu plates. For the *bur1*Δ bypass assay, dilutions were spotted onto SC–Leu–Ura plates with and without 5-FOA to select against the pRS316-Bur1 plasmid. Growth was assayed after between 48 and 96 hours as indicated for plates placed either in the dark or in 500 *μ*W/cm^2^ blue light emitted from an LED strip with maximum emission at 465 nm.

### Steady-state immunoblotting

BY4741 wild type and *set2*Δ overnight cultures were transformed and plated on SC–Leu plates that were then incubated in the dark at 30 °C. A colony from the LANS-Set2 transformation was resuspended in SC–Leu and split into light and dark cultures, whereas all other transformants were resuspended and grown in the dark. A colony from the Set2-FRB strain was resuspended in YPD and split into cultures either lacking rapamycin or with exposure to 1 *μ*g/mL rapamycin in ethanol. All cultures were placed in the same incubator overnight at 30 °C: dark cultures were wrapped in foil and light cultures were exposed to 500 *μ*W/cm^2^ blue light (465 nm) from an LED strip wrapped around the base of the tube rack. In the morning cell density was measured at OD_600_ and cultures were diluted in SC–Leu to a final OD_600_ of 0.3 in a final volume of 6.5 mL. Cultures were then returned to an incubator at 30 °C in either the same light conditions or the same rapamycin exposure conditions for 5 hours, after which OD600 was measured and 5 OD600 units of each asynchronous log phase culture were collected. Samples for chemiluminescent detection (Figures S1C and S1E and all Set2 and G6PDH blots) were processed as follows: cells were collected by centrifugation and lysed using glass beads and vortexing at 4 °C for 8 minutes in SUMEB (1% SDS, 8 M urea, 10 mM MOPS, pH 6.8, 10 mM EDTA, 0.0 % bromophenol blue). Extracts were retrieved, centrifuged and boiled at 95 °C for 5 minutes. Samples for near-infrared fluorescent detection (all other figures) were processed as follows: cells were added to the appropriate volume of 100% TCA (Sigma, 100% w/v) to obtain a final concentration of 20% TCA, mixed and centrifuged at 5k RPM. Supernatants were discarded and pellets were stored at –80 °C. After freezing, samples were resuspended in TCA buffer (10 mM Tris, pH 8.0, 10% TCA, 25 mM NH4OAc, 1 mM Na2 EDTA), mixed and incubated for 10 minutes on ice. Samples were centrifuged, resuspended in resuspension buffer (0.1 M Tris, pH 11.0, 3% SDS), and boiled at 95 °C for 10 minutes. Samples were centrifuged to clarify the extracts, protein was quantified using the DC assay and samples were diluted in resuspension buffer to equivalent concentrations and further diluted in 2× SDS Loading Buffer (60 mM Tris pH 6.8, 2% SDS, 10% glycerol, 0.2% bromophenol blue, 100 mM DTT added fresh). 10 *μ*L of whole cell extracts from either preparation method were loaded on 15% SDS-PAGE gels (or 8% SDS-PAGE for the Set2 blot). Proteins were transferred to 0.45 *μ*m PVDF membranes (Millipore Sigma Immobilon-FL for near-infrared fluorescent detection) using a Hoefer Semi-Dry Transfer Apparatus at 45 mA per membrane. For chemiluminescent blotting, primary antibodies were incubated in 5% milk at 4 °C overnight and secondary antibodies were incubated in 5% milk for 1 hour. Immunoblots were developed using ECL Prime (Amersham RPN2232). For near-infrared fluorescent blotting, primary antibodies were incubated in Odyssey blocking buffer at 4 °C overnight and secondary antibodies were incubated in Odyssey blocking buffer with 0.015% SDS for 1 hour.

### Photoactivation immunoblotting

Colonies from LANS-Set2 transformations of the appropriate yeast strain were resuspended in SC–Leu and grown in the dark or 500 *μ*W/cm^2^ blue light overnight at 30 °C. For the Set2-FRB anchor away strain colonies were resuspended in YPD and grown without rapamycin exposure at 30 °C. In the morning cell density was measured at OD_600_ and cultures were diluted in SC–Leu to a final volume of 70 mL and final OD_600_ of 0.35 (OD_600_ of 0.3 for Set2-FRB). For *set2*Δ*bar1*Δ cultures, after 3 hours cultures were split in two cultures, one of which was treated with 0.1 *μ*g/mL of alpha-factor (Diag #RP01002), grown another 1.5 hours, visualized to ensure arrest in treated cells, and grown another 30 minutes. All other cultures were grown in the same light conditions, though light cultures were grown in 200 *μ*W/cm^2^ blue light (465 nm emitted from an LED strip above the culture flasks) for 4.5 hours of growth at 30 °C at which point 5 OD_600_ units of each asynchronous log phase culture were collected. Time courses began when cultures were moved from the dark to light or the light to dark (or for the Set2-FRB strain upon addition of 1 *μ*g/mL rapamycin in ethanol). At each time point the same volume of each culture (5 OD_600_ units measured at time zero) was harvested by the TCA method detailed above. Frozen pellets were further processed as above after the completion of each time course.

### Immunoblot quantification

Band intensities for H3, H3K36me3, H3K36me2, H3K36me1, and H3K56ac were quantified with local background subtracted using Image Studio Lite. H3K36 methylation and H3K56 acetylation intensities were divided by their respective H3 intensities. To obtain relative abundance throughout the dark to light time courses, the last time point was used as a reference for each preceding time point, and for the light to dark and Anchor Away time courses the first time point was used as a reference for each subsequent time point. Relative abundances were plotted using GraphPad Prism 5, statistical significance was calculated using unpaired two-tailed student’s *t*-test (*p* < 0.05), and half-lives were obtained by fitting data using single exponentials. The one phase association equation *Y* = *Y*_0_ + (plateau – *Y*_0_)(l – *e^−kx^*) was used to fit values corresponding to the LANS-Set2 dark-to-light time courses whereas the one phase decay equation *Y* = (*Y*_0_ – plateau)*e^−kx^*) + plateau was used to fit values corresponding to the LANS-Set2 light-to-dark as well as the Anchor Away time courses.

### Chromatin immunoprecipitation (ChIP)

For WT, *set2*Δ and *rph12*Δ strains, yeast was resuspended in YPD and grown in the dark overnight at 30 °C. For time courses, colonies from LANS-Set2 transformations of the appropriate yeast strain were resuspended in SC–Leu and grown in the dark or 500 *μ*W/cm^2^ blue light overnight at 30 °C. In the morning cell density was measured at OD600 and cultures were diluted in appropriate media to a final volume of 70 mL and final OD_600_ of 0.35. Cultures were maintained in the dark or light (200 *μ*W/cm^2^ blue light) for 4.5 hours of growth at 30 °C until the OD_600_ reached 0.8-1.0. For WT, *set2*Δ and *rph12*Δ strains samples were collected, and for time courses an initial time point was collected then samples were shifted from the dark to light or the light to dark. To collect samples, cells were fixed with a 1% final concentration of formaldehyde, the fixation was quenched, cells were washed, and pellets were frozen at −80 °C. After the completion of each time course, cells were lysed using 140 mM FA lysis buffer (containing protease inhibitor cocktail). Lysed cells were sonicated (Diagenode Bioruptor UCD-200) on high intensity 5 times for 5 minutes each (cycles of 30 seconds “ON” and 30 seconds “OFF”) and clarified by centrifuging at full speed for 15 minutes. Overnight immunoprecipitations of *S. cerevisiae* chromatin with appropriate antibodies were prepared with *S. pombe* chromatin spike-in control corresponding to 15% of *S. cerevisiae* chromatin as estimated by Bradford assay (Bio-Rad). 50 *μ*L of Protein G Dynabeads were added to each 500 *μ*L immunoprecipitation reaction and reactions were incubated for 2 hours. Washes were performed with 1 mL of 140 mM FA-lysis buffer, 500 mM FA-lysis buffer, LiCl solution (250 mM LiCl, 10 mM Tris, 0.5% each of NP-40 and sodium doxycholate and 1 mM EDTA) and TE pH 8.0. Elution buffer (1% SDS, 0.1 M NaHCO_3_) was used to elute the DNA (15 minutes shaking at 65 °C followed by centrifugation at 2,000 RPM for 2 minutes). 10 *μ*L of 5 M NaCl was added to the eluates and 10% inputs and samples were incubated at 65 °C overnight to carry out de-crosslinking. Samples were treated with RNase for 1 hour and proteinase K for 1 hour before ChIP DNA Clean & Concentrator (Zymo Research) to extract the DNA.

### ChIP-sequencing and data analysis

Bar-coded sequencing libraries were prepared as recommended by the manufacturer (KAPA Hyper Prep Kit), pooled and sequenced (Hi-Seq 2500, Illumina). Reads from the sequencer were demultiplexed using bcl2fastq (v2.20.0). Sequencing adapters on reads were trimmed using cutadapt (v1.12) using options -a GATCGGAAGAGC -A GATCGGAAGAGC and `--minimum-length 36` in paired mode. After trimming, reads were filtered for quality using the `fastq_quality_filter` in FASTX-Toolkit (v0.0.12), with options `-Q 33, -p 90`, and `-q 20`. In-house scripts were used to limit potential PCR duplicates by limiting reads with the same sequence to a maximum of 5 copies and discarding the copies beyond that limit. Since the previous filtering steps may remove one end of a read pair but not the other, both read-pair fastq files were synced using in-house scripts to ensure proper order for alignment. As the ChIP experiments contained *S. pombe* spike-in, a chimeric *S. cerevisae-S. pombe* genome was generated using the genomeGenerate tool in STAR (v2.5.2b). This chimeric genome contains the full sequence of both species, allowing reads to align to their best overall fit between the two species. Once generated, read alignment was done using STAR (v2.5.2b) and options `--outFilterMismatchNmax 2, --chimSegmentMin 15, --chimJunctionOverhangMin 15, --outSAMtype BAM Unsorted, --outFilterType BySJout, --outFilterScoreMin 1`, and `--outFilterMultimapNmax 1`. Post-alignment, Samtools (v1.31) and bedtools (2.26) were used to generate bigWig files for downstream analyses. To account for *S. pombe* spike-in, the bigwig signal was normalized by the total number of *S. pombe* reads per million, using the -scale option within the bedtools genomecov tool. Normalized H3K36me3 signal was obtained by dividing H3K36me3 spike-in normalized signal by H3 spike-in normalized signal for each base pair for each replicate. Some regions were devoid of any H3 signal, and these regions were flagged and excluded from further analyses.

Deeptools (v2.5.4) was used to generate metagene plots using the normalized H3K36me3 signal. Deeptools was also used to obtain base pair by base pair signal over regions of interest such as genes, introns, and exons. This data was used to calculate the average signal per gene, excluding those regions flagged for lacking H3 signal. We set out to understand H3K36me3 signals over time, however differing levels of normalized H3K36me3 signal within genes made analysis difficult. To account for these differences, we scaled the average signal per gene throughout the time course between 0 (no signal) and 1 (the maximum signal of a gene over the time course). This yielded a relative scale for each gene as a fraction of its maximum signal, and allowed for easier comparisons of patterns within and across treatments.

### RNA isolation

Colonies from LANS-Set2 transformations in *set2*Δ were prepared as above, except that in the morning cultures were diluted in SC–Leu to a final volume of 70 mL and final OD_600_ of 0.3. Time courses were conducted as above except that 10 mL of log-phase cultures were collected by centrifugation and frozen at −80 °C. After the completion of each time course, RNA was isolated by acid phenol extraction. RNA (10 *μ*g) was treated with DNase (Promega) and purified (RNeasy column, QIAGEN).

### RNA-sequencing and data analysis

RNA (2.5 *μ*g) was processed using rRNA depletion beads specific to yeast (Illumina). Bar-coded sequencing libraries were prepared as recommended by the manufacturer (TruSeq Stranded Total RNA Library Preparation Kit, Illumina), pooled and sequenced (Hi-Seq 4000, Illumina). Reads from the sequencer were demultiplexed using bcl2fastq (v2.20.0). Reads were trimmed using cutadapt (v1.12) using options - a GATCGGAAGAGC -A GATCGGAAGAGC and `--minimum-length 36` to remove any sequencing adapters. After trimming, reads were filtered for quality using the `fastq_quality_filter` in FASTX-Toolkit (v0.0.12), with options `-Q 33, -p 90`, and `-q 20`. Reads were aligned to the sacCer3 genome using STAR (v2.5.4b) with options --quantMode TranscriptomeSAM, --outFilterMismatchNmax 2, --alignIntronMax 1000000, --alignIntronMin 20, --chimSegmentMin 15, --chimJunctionOverhangMin 15, --outSAMtype BAM Unsorted, --outFilterType BySJout, and --outFilterMultimapNmax 1. To calculate the RNA abundance values, Salmon (v0.8.1) tool was used with options `-l SR, -- incompatPrior 0.0` to account for read strandedness. Samtools (v1.3.1), bedtools (v2.26), and R (v3.3.1) were used to interconvert files for downstream analyses. DESeq2 (v1.14.1) was used to determine which genes had differential expression. Venn diagrams were made using R package Vennerable (v3.0).

### Bayesian generalized linear mixed effect model

A Bayesian generalized linear mixed effect model, implemented in R (v3.5.2) with the brms (v2.8.0) and rstan (v2.18.2) packages as wrappers for the statistical software Stan (v2.18.1), to account for features of the data: non-normality and the longitudinal and replicate observation study design. Posterior summaries from the models were used to make inferences and for further analysis. See Supplemental Methods for greater detail.

## DATA ACCESS

All raw and processed sequencing data generated in this study have been submitted to the NCBI Gene Expression Omnibus (GEO; https://www.ncbi.nlm.nih.gov/geo/) under accession number GSE123456.

## ACKNOWLEDGMENT

This work was supported by the NIH (R01DA036877, R35GM126900, and F31GM122321).

## Author contributions

Conceptualization, A.M.L., A.J.H., B.D.S, B.K., I.J.D; Methodology, A.M.L., A.J.H., G.R.K., H.Y.; Software, A.J.H., G.R.K.; Formal Analysis, A.J.H., G.R.K.; Investigation, A.M.L., H.M., H.Y., D.R., S.Z.; Resources, J.B.; Data Curation, A.J.H., G.R.K.; Writing – Original Draft, A.M.L., A.J.H., B.D.S., I.J.D.; Writing – Review & Editing, all authors; Visualization, A.M.L., A.J.H., G.R.K.; Supervision, I.J.D., B.K., B.D.S.; Funding Acquisition, A.M.L., H.Y., B.K., B.D.S., I.J.D.

