## Supplemental Material for "An optogenetic switch for the Set2 methyltransferase provides evidence for rapid transcription-dependent and independent dynamics of H3K36 methylation"

**3 Supplemental Movie** **13**

**4 Supplemental Note** **13**

**5 Supplemental Statistical Methods** **14**

### 1 Supplemental Figures

#### 1.1 Figure S1: Related to Figures 1 and 2.

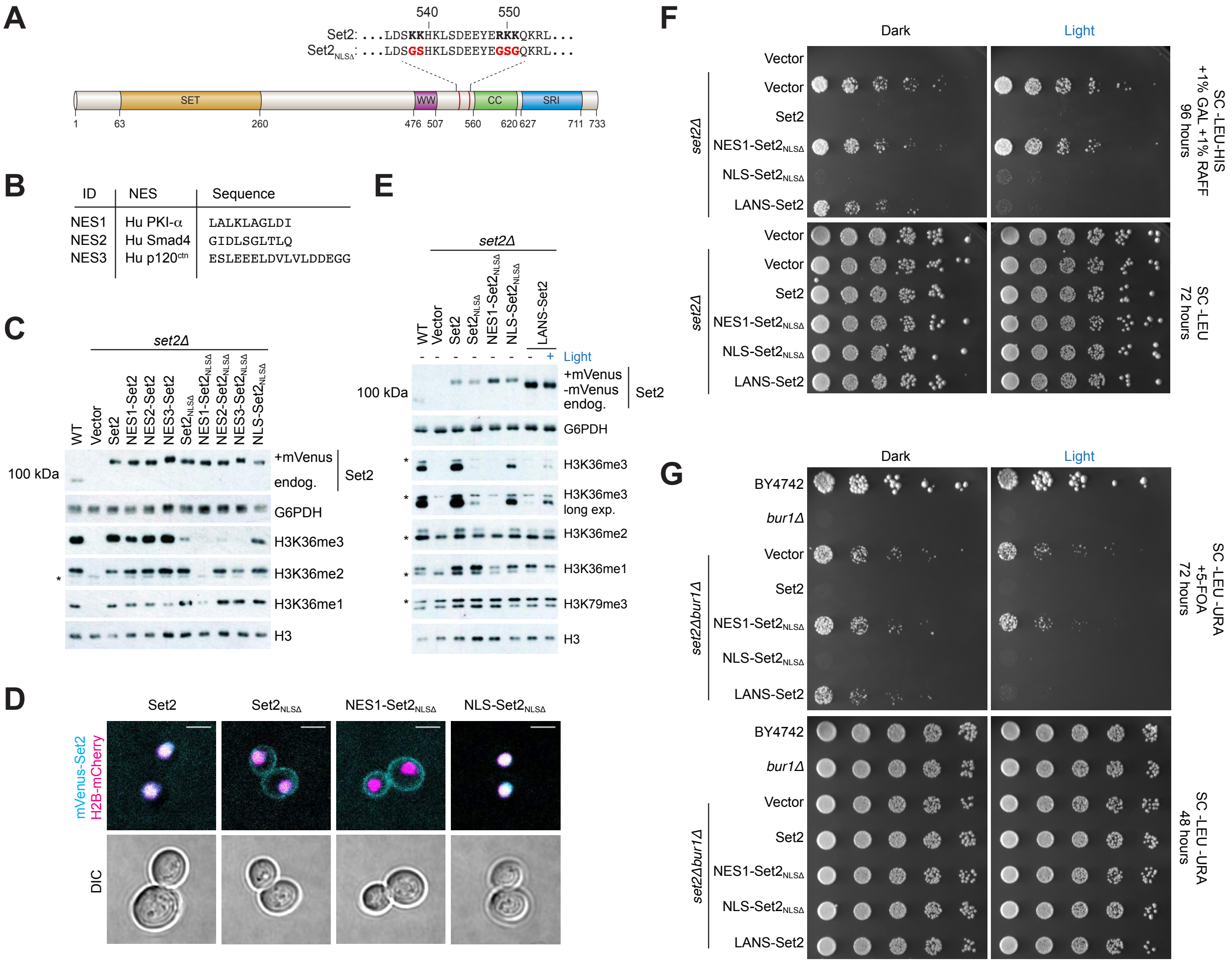

(A) Sequence alignment and domain map depicting mutations (red) to a putative NLS (bold) and their locations relative to other domains. Set2 is characterized by the SET (**S**u(var)3-9, **E**nhancer-of-zeste and **T**rithorax), WW, CC (coiled coil), and SRI (Set2-Rpb1 interacting) domains. (B) List of the nuclear export signals tested, including their origins and sequences. (C) Immunoblots showing testing in a *SET2* deletion strain (*set2*Δ) of nuclear export signals fused to Set2 with and without inactivating mutations of the Set2 NLS ($\text{Set2}_{\text{NLS}\Delta}$), as well as testing of the designed NLS used in LANS. Constructs are mVenus-tagged and migrate around 115 kDa, whereas the endogenous Set2 is not mVenus-tagged and migrates around 85 kDa. The asterisk indicates a nonspecific band. (D) Confocal images demonstrating localization of several mVenus-tagged Set2 constructs (cyan) in yeast cells with histone H2B endogenously tagged with mCherry (magenta); (scale bar, 3 $\mu$m). (E) Light-mediated control of H3K36 tri- and dimethylation in *set2*Δ cells as evidenced by immunoblotting with antibodies specific to various histone modifications. Several constructs are mVenus-tagged whereas endogenous Set2 (endog.) and LANS-Set2 are not. Asterisks indicate nonspecific bands. (F) *FLO8-HIS3* cryptic transcription initiation spotting assay testing LANS-Set2 and constructs toward its development. Six-fold serial dilutions of overnight cultures expressing one of several constructs were spotted on solid media with or without histidine as per the *FLO8-HIS3* reporter assay depicted in

Figure 2C, and plates were incubated in the dark or blue light. Images were collected 3-4 days later. All constructs are mVenus-tagged. (G) *bur1*Δ bypass spotting assay testing LANS-Set2 and constructs toward its development. Five-fold serial dilutions of overnight cultures of wild-type *BY4742* and *BUR1* plasmid shuffling strains were spotted on solid media with or without 5-FOA to select against the *BUR1*/*URA3* plasmid and plates were incubated in the dark or blue light. Images were collected 2-3 days later. All constructs are mVenus-tagged.

#### 1.2 Figure S2: Related to Figure 2.

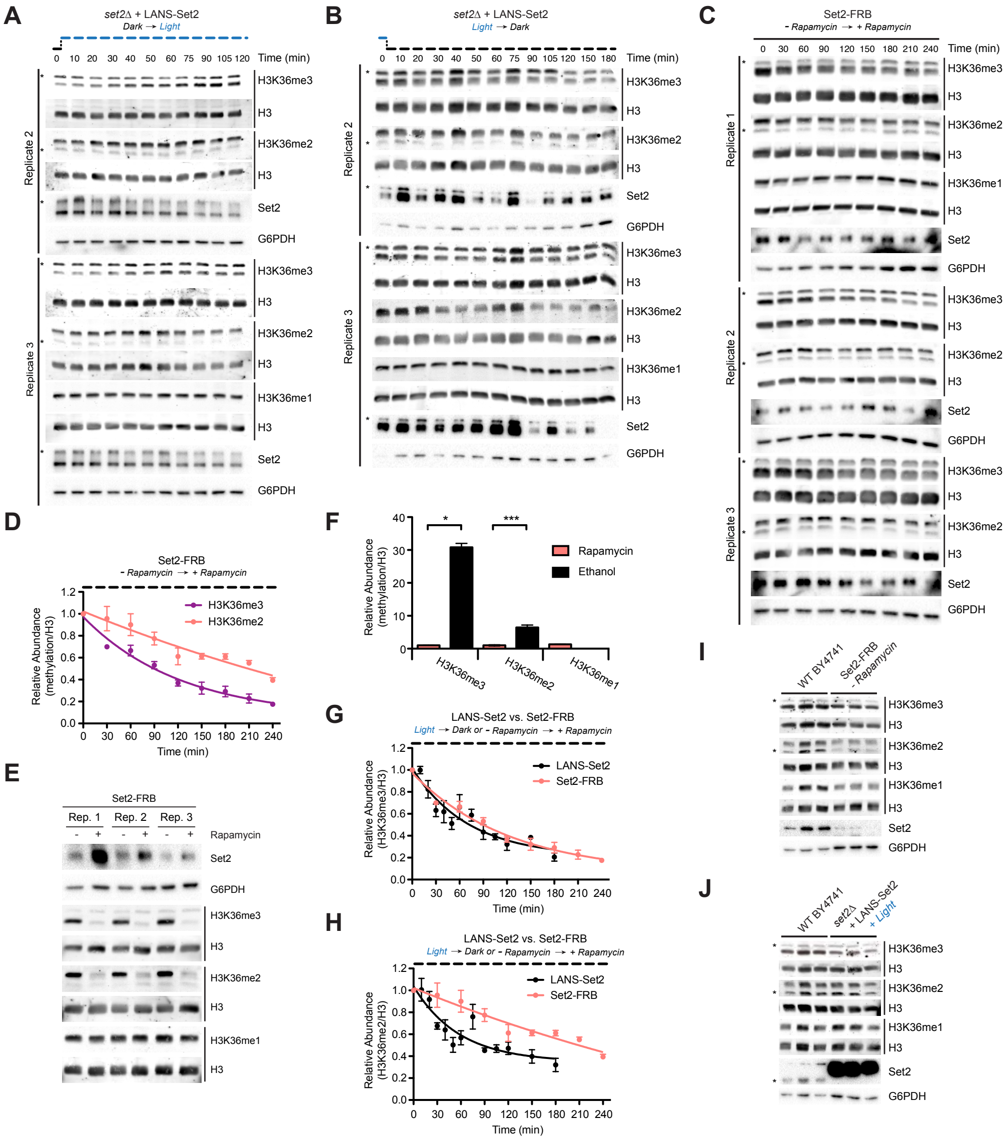

(A) Western blots probing gain of H3K36 methylation over time using LANS-Set2 in *set2*Δ after the transition from dark to light. Triplicate immunoblots of whole-cell lysates prepared from log phase cultures for quantification of histone modifications as a function of time in Figure 2F. Asterisks indicate nonspecific bands. (B) Western blots probing loss of H3K36 methylation over time using LANS-Set2 in *set2*Δ after the transition from light to dark. Triplicate immunoblots of whole-cell lysates prepared from log phase cultures for quantification of histone modifications as a function of time in Figure 2H. Asterisks indicate nonspecific bands. (C) Western blots probing loss of H3K36 methylation over time using Set2-FRB after addition of rapamycin. Triplicate immunoblots of whole-cell lysates prepared from log phase cultures. (D) Quantification of histone modifications as a function of time from immunoblots in (C). ($n$ = 3, mean ± SEM). (E) Immunoblots comparing levels of H3K36 methylation in Set2-FRB cultures grown with and without rapamycin exposure. (F) Quantification of histone modifications from immunoblots in (E). Data represent mean values ± SD ($n$ = 3 independent experiments). (G) Quantification of H3K36me3 loss using LANS-Set2 in *set2*Δ after transition from light to dark (from Figure 2F) and using Set2-FRB after addition of rapamycin (from

Figure S2D). (H) Quantification of H3K36me2 loss using LANS-Set2 in *set2*Δ after transition from light to dark (from Figure 2H) and using Set2-FRB after addition of rapamycin (from Figure S2D). (I) Triplicate immunoblots of whole-cell lysates prepared from log phase cultures for wild-type BY4741 yeast and Set2-FRB without exposure to rapamycin. Asterisks indicate nonspecific bands. (J) Triplicate immunoblots of whole-cell lysates prepared from log phase cultures for wild-type BY4741 yeast and LANS-Set2 in *set2*Δ grown continuously in the light. Asterisks indicate nonspecific bands. Half-lives were calculated from single exponential fits to the H3K36me3 and H3K36me2 relative abundance data using GraphPad Prism 5. $*P < 0.05; ***P < 0.0001$.

#### 1.3 Figure S3: Related to Figure 3.

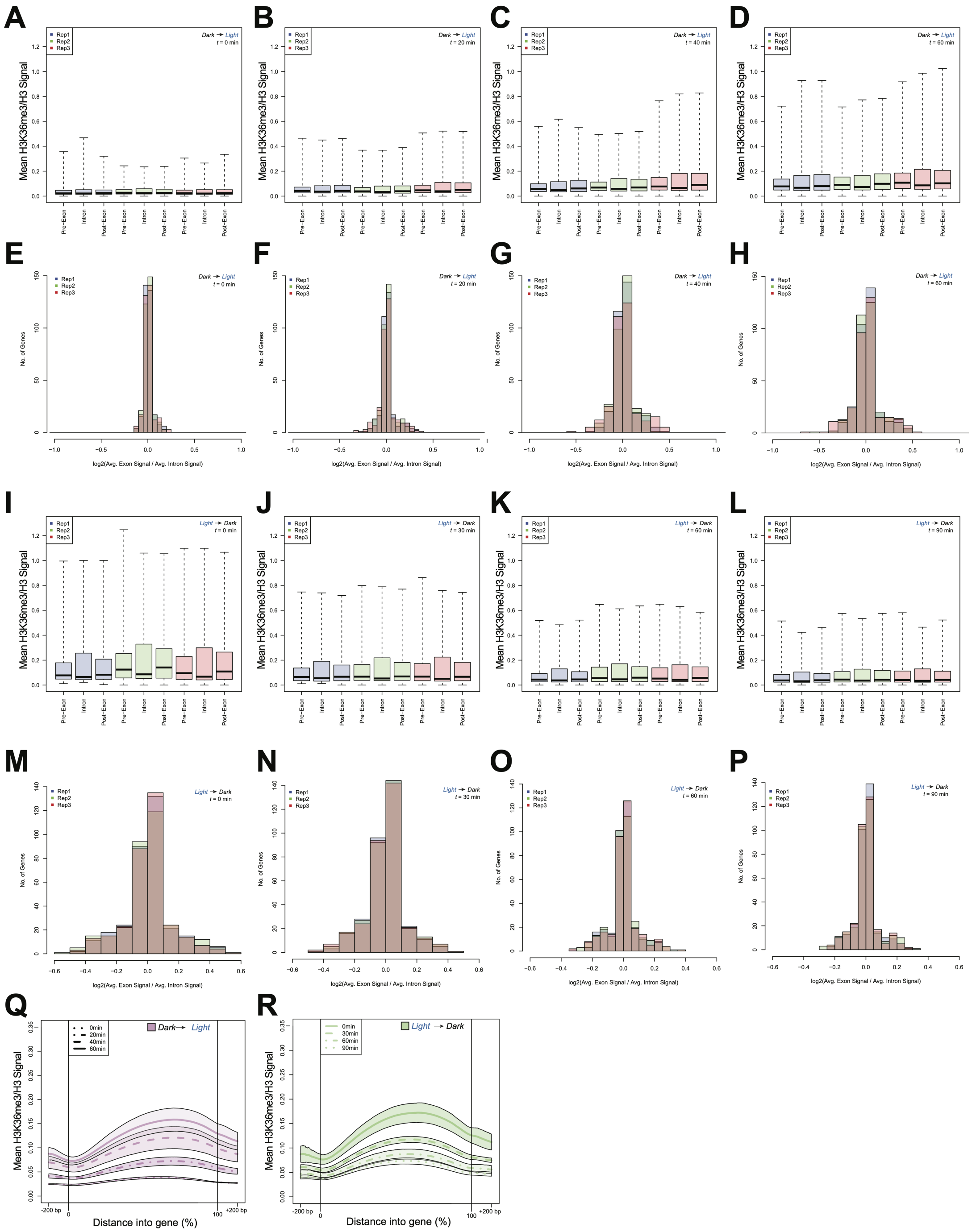

(A,B,C,D) Boxplots representing the distribution of all mean, normalized H3K36me3 ChIP-seq signal for the pre-exons, introns, and post-exons at (A) $t = 0$ minutes, (B) $t = 20$ minutes, (C) $t = 40$ minutes, and (D) $t = 60$ minutes after LANS-Set2 activation. A pre-exon is the exon preceding a given intron while the post-exon is the succeeding exon to a given intron. Blue boxplots are replicate 1, green boxplots are replicate 2, and blue boxplots are replicate 3. Boxplot borders represent the $2^{nd}$ and $3^{rd}$ quartile, while the midline reflects the median. (E,F,G,H) Histograms representing the $\log_2$ fold change of mean, normalized

H3K36me3 ChIP-seq signal between a pre-exon and the corresponding intron at (E) $t = 0$ minutes, (F) $t = 20$ minutes, (G) $t = 40$ minutes, and (H) $t = 60$ minutes after LANS-Set2 activation. Blue histograms represent replicate 1, green histograms represent replicate 2, and blue histograms represent replicate 3. (I,J,K,L) Boxplots representing the distribution of all mean, normalized H3K36me3 ChIP-seq signal for the pre-exons, introns, and post-exons (I) $t = 0$ minutes, (J) $t = 30$ minutes, (K) $t = 60$ minutes, and (L) $t = 90$ minutes after LANS-Set2 inactivation. (M,N,O,P) Histograms representing the $\log_2$ fold change of mean, normalized H3K36me3 ChIP-seq signal between a pre-exon and the corresponding intron at (M) $t = 0$ minutes, (N) $t = 30$ minutes, (O) $t = 60$ minutes, and (P) $t = 90$ minutes after LANS-Set2 inactivation. (Q,R) Mean, normalized H3K36me3 ChIP-seq signal metagene for the time course after (Q) LANS-Set2 activation or (R) LANS-Set2 inactivation, scaled between 0-100% of the gene length. Metagenes contain 200 base pairs up- and downstream of the genes. Bold lines represent the mean across all three replicates for each time point, while the shaded bands represent the standard deviation about the mean. The different line types indicate the time points.

#### 1.4 Figure S4: Related to Figure 4.

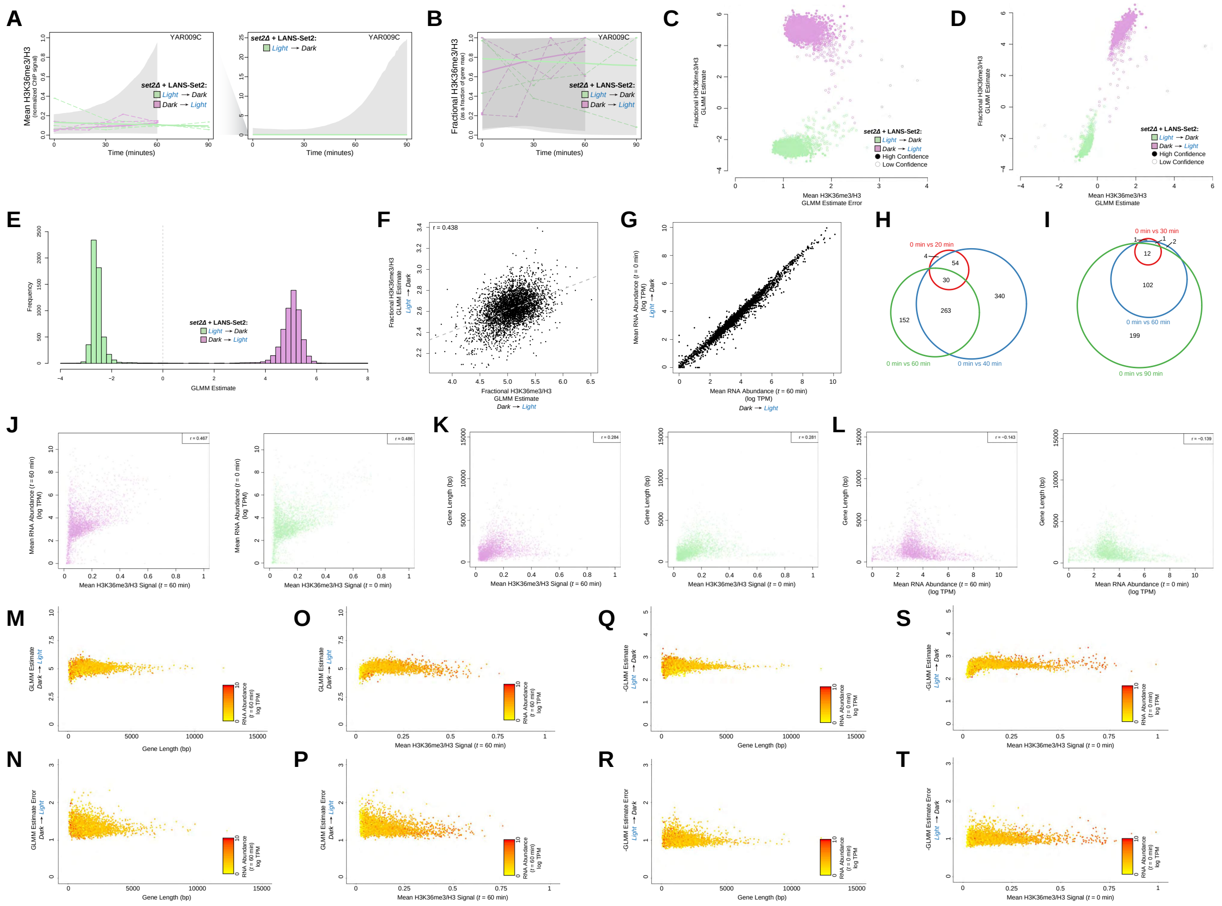

The (A) mean, normalized H3K36me3 ChIP-seq signal and (B) relative H3K36me3 ChIP-seq signal for the gene YAR009C throughout the time course after LANS-Set2 activation (green) and LANS-Set2 inactivation (purple). Dashed lines represent individual replicates, while bold lines represent the Bayesian generalized linear mixed effect model (GLMM) posterior mean H3K36me3. The shaded regions represent the posterior credible interval for H3K36me3 from the GLMM. YAR009C was not included in the high confidence gene set from Figure **??**B, which is reflected in the lack of a clear trend in the data and wide credible intervals for either normalized or relative H3K36me3. The credible interval for LANS-Set2 activation was plotted on a separate plot (right) to accommodate the large difference in the y-axis scale. (C) Per-gene GLMM posterior rates for relative H3K36me3 ChIP-seq signal for LANS-Set2 activation (green) and LANS-Set2 inactivation (purple) and their respective errors. Solid circles represent high confidence genes for LANS-Set2 activation or LANS-Set2 inactivation, while hollow circles represent genes with low confidence. (D) Per-gene GLMM posterior rates for normalized H3K36me3 ChIP-seq signal and relative H3K36me3 ChIP-seq signal for both LANS-Set2 activation (green) and LANS-Set2 inactivation (purple). All genes were included, whereas some were excluded in Figure 4C for clarity. (E) Histogram of the GLMM rates for of all genes for LANS-Set2 activation (green) and LANS-Set2 inactivation (purple). (F) GLMM rate comparison between LANS-Set2 inactivation and LANS-Set2 activation for the high confidence gene set. Pearson correlation coefficient is included. (G) Comparison of mean RNA abundance levels (log TPM) between LANS-Set2 inactivation and LANS-Set2 activation at $t = 0$ minutes and $t = 60$ minutes, respectively. (H) Venn diagram of genes with significantly different RNA abundances between $t = 0$ minutes and $t = 30$ minutes (red), $t = 60$

minutes (blue), and $t = 90$ minutes (green) after LANS-Set2 activation. (I) Venn diagram of differential genes between $t = 0$ minutes and $t = 20$ minutes (red), $t = 40$ minutes (blue), and $t = 60$ minutes (green) after LANS-Set2 inactivation. (J) Mean RNA abundance levels (log TPM) by mean, normalized H3K36me3 levels for the high confidence gene set for both LANS-Set2 activation at $t = 60$ minutes (left) and LANS-Set2 inactivation at $t = 0$ minutes (right). These time points were chosen as they are the ones with the highest H3K36me3 level, and thus most like wild-type conditions. (K) Gene length by mean, normalized H3K36me3 levels for the high confidence gene set for both LANS-Set2 activation at $t = 60$ minutes (left) and LANS-Set2 inactivation at $t = 0$ minutes (right). (L) Gene length by mean RNA abundance levels (log TPM) for the high confidence gene set for both LANS-Set2 activation at $t = 60$ minutes (left) and LANS-Set2 inactivation at $t = 0$ minutes (right). (M) The GLMM rate by gene length for the high confidence gene set during LANS-Set2 activation. (N) GLMM rate errors by gene length for the high confidence gene set during LANS-Set2 activation. (O) The GLMM rate by mean, normalized H3K36me3 levels for the high confidence gene set during LANS-Set2 activation. (P) The GLMM rate errors by mean, normalized H3K36me3 levels for the high confidence gene set during LANS-Set2 activation. Genes are colored based on their RNA abundance at $t = 60$ minutes (log TPM) (M,N,O,P). (Q) The GLMM rate by gene length for the high confidence gene set during LANS-Set2 inactivation. (R) GLMM rate errors by gene length for the high confidence gene set during LANS-Set2 inactivation. (S) The GLMM rate by mean, normalized H3K36me3 levels for the high confidence gene set during LANS-Set2 inactivation. (T) The GLMM rate errors by mean, normalized H3K36me3 levels for the high confidence gene set during LANS-Set2 inactivation. Genes are colored based on their RNA abundance at $t = 0$ minutes (log TPM) (Q,R,S,T).

#### 1.5 Figure S5: Related to Figure 5.

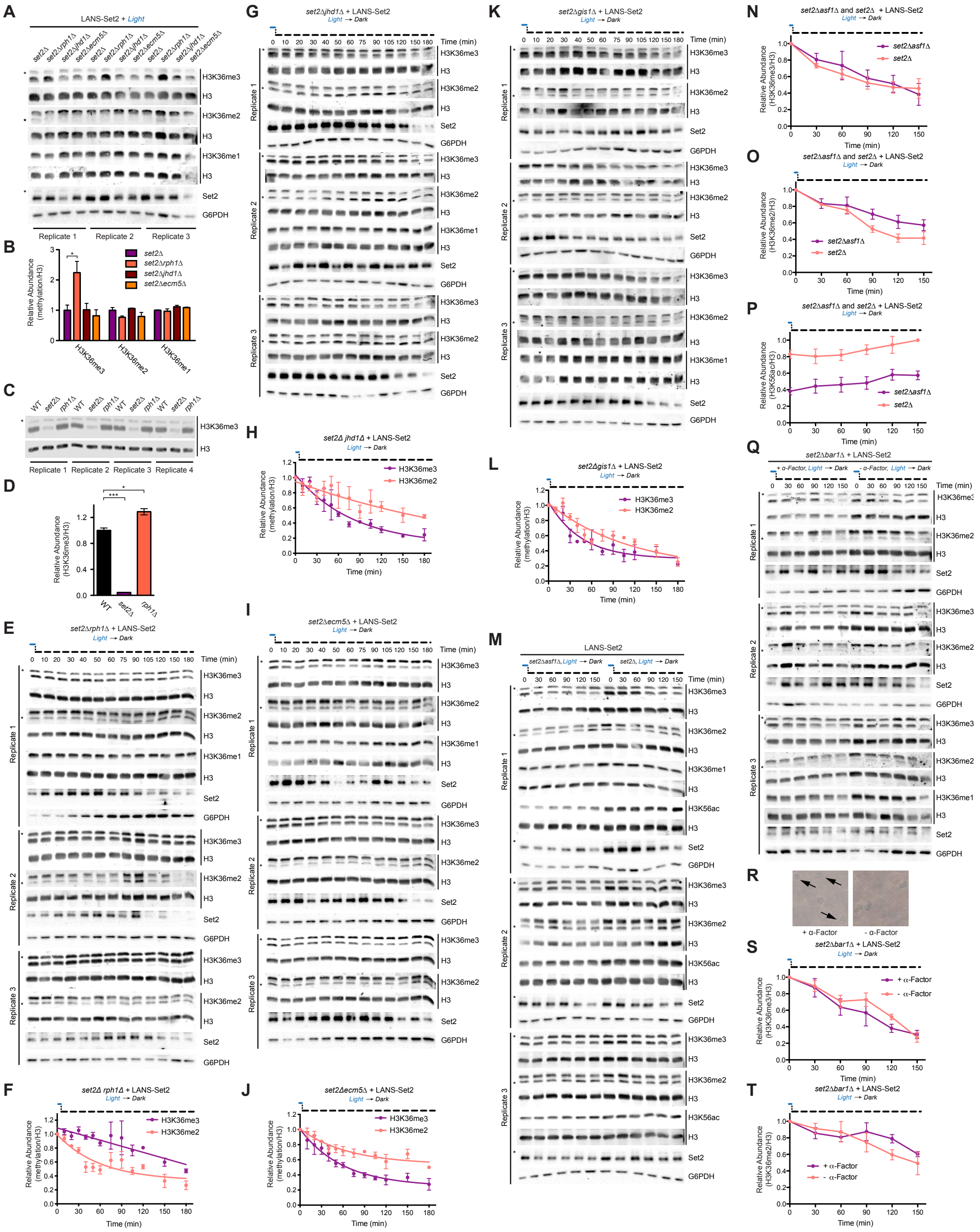

(A) Triplicate immunoblots of whole-cell lysates prepared from log phase cultures for the indicated strains transformed with LANS-Set2 and grown continuously in the light. Asterisks indicate nonspecific bands. (B) Quantification of Western blots in (A). Data represent mean values ± SD ($n = 3$). (C) Replicate immunoblots of whole-cell lysates prepared from log phase cultures for the indicated strains. (D) Quantification of Western blots in (C). Data represent mean values ± SD ($n = 4$). (E) Western blots probing loss of H3K36 methylation over time using LANS-Set2 in *set2*Δ*rph1*Δ. Triplicate immunoblots of whole-cell lysates prepared from log phase cultures for quantification of histone modifications as a function of time after the transition from the light to dark. Asterisks indicate nonspecific bands. (F) Quantification of immunoblots in (E); ($n = 3$, mean ± SEM). (G) Western blots probing loss of H3K36 methylation over time using LANS-Set2 in *set2*Δ*jhd1*Δ. Triplicate immunoblots of whole-cell lysates prepared from log phase cultures for quantification of histone modifications as a function of time after the transition from the light to dark. Asterisks indicate nonspecific bands. (H) Quantification of immunoblots in (G); ($n = 3$, mean ± SEM). (I) Western blots probing loss of H3K36 methylation over time using LANS-Set2 in *set2*Δ*ecm5*Δ. Triplicate immunoblots of whole-cell lysates prepared from log phase cultures for quantification of histone modifications as a function of time after the transition from the light to dark. Asterisks indicate nonspecific bands. (J) Quantification of immunoblots in (I); ($n = 3$, mean ± SEM). (K) Western blots probing loss of H3K36 methylation over time using LANS-Set2 in *set2*Δ*gis1*Δ. Triplicate immunoblots of whole-cell lysates prepared from log phase cultures for quantification of histone modifications as a function of time after the transition from the light to dark. Asterisks indicate nonspecific bands. (L) Quantification of immunoblots in (K); ($n = 3$, mean ± SEM). (M) Western blots probing loss of H3K36 methylation over time using LANS-Set2 in *set2*Δ and *set2*Δ*asf1*Δ. Triplicate immunoblots of whole-cell lysates prepared from log phase cultures after the transition from the light to dark. Asterisks indicate nonspecific bands. (N) Quantification of H3K36me3 from immunoblots in (M); ($n = 3$, mean ± SEM). (O) Quantification of H3K36me2 from immunoblots in (M); ($n = 3$, mean ± SEM). (P) Quantification of H3K56ac from immunoblots in (M); ($n = 3$, mean ± SEM). (Q) Western blots probing loss of H3K36 methylation over time with and without $\alpha$-factor treatment using LANS-Set2 in *set2*Δ*bar1*Δ. Triplicate immunoblots of whole-cell lysates prepared from log phase cultures grown with and without pre-treatment with $\alpha$-factor to arrest cells in G1 prior to transition from the light to dark. Asterisks indicate nonspecific bands. (R) Representative widefield microscopy images of *set2*Δ*bar1*Δ cells with and without $\alpha$-factor pre-treatment. Arrows point to the aberrant morphology (“shmoo" shape) that develops after $\alpha$-factor treatment of MATa cells. (S) Quantification of H3K36me3 from immunoblots in (Q); ($n = 3$, mean ± SEM). (T) Quantification of H3K36me2 from immunoblots in (Q); ($n = 3$, mean ± SEM). Half-lives were calculated from single exponential fits to the H3K36me3 and H3K36me2 relative abundance data using GraphPad Prism 5. $*P < 0.05$; $***P < 0.0001$.

#### 1.6 Figure S6: Related to Figure 6.

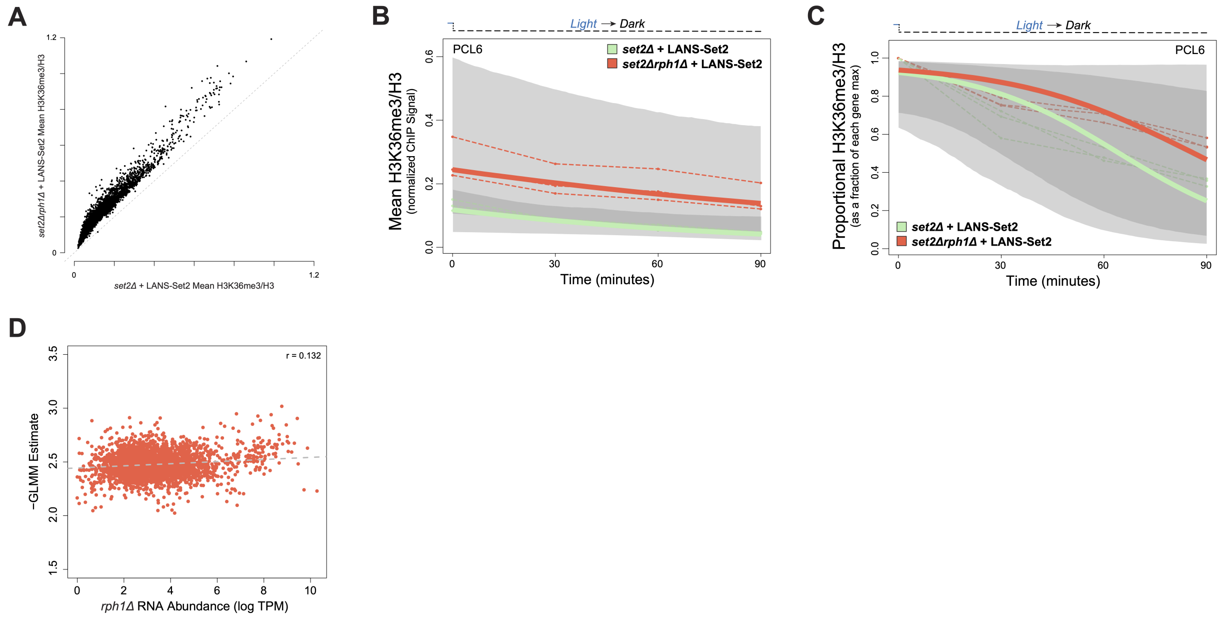

(A) Comparison of mean, normalized H3K36me3 ChIP-seq signal for all genes after LANS-Set2 inactivation between *set2*Δ and *set2*Δ*rph1*Δ backgrounds. The (B) normalized H3K36me3 ChIP-seq signal and (C) relative H3K36me3 ChIP-seq signal for the gene *PCL6* (YAL012W) throughout the time courses of LANS-Set2 inactivation in the backgrounds of *set2*Δ (green) or *set2*Δ*rph1*Δ (red). Dashed lines represent individual replicates, while bold lines represent the posterior mean H3K36me3 from the Bayesian generalized linear mixed effect model (GLMM). The shaded regions represent the posterior credible interval for H3K36me3 from the GLMM. (D) Comparison of the LANS-Set2 inactivation GLMM rate in *set2*Δ*rph1*Δ cells and RNA abundance levels (log TPM) in *rph12*Δ cells across the high confidence gene set. Dashed line represents the line of best fit. Pearson correlation coefficient is $r = 0.131$.

### 2 Supplemental Tables

#### 2.1 Table S1: Yeast strains used in this study.

| Name | Genotype | Source |
|---|---|---|
| BY4741 | MATa, *his3Δ0 leu2Δ0 met15Δ0 ura3Δ0* | Open Biosystems, Huntsville, Alabama |
| *set2Δ* | MATa, *his3Δ0 leu2Δ0 met15Δ0 ura3Δ0 set2Δ*::HPH | this study |
| H2B-mCherry | MATα *trp1Δ63 leu2Δ1 ura3-52 his3Δ200 lys2-801* H2B-mCherry::NatMX | Gift of Kerry Bloom |
| KLY78 | MATx, *ura3-52 leu2Δ1 or leu1- trp1Δ63 or trp1- his3Δ200* KanMX6::GAL1pr-flo8::HIS3 *lys2-128Δ* | Silva et al, J. Biol. Chem., 2012 |
| KLY78 *set2Δ* (YSM138) | MATx, *ura3-52 leu2Δ1 or leu1- trp1Δ63 or trp1- his3Δ200* KanMX6::GAL1pr-flo8::HIS3 *lys2-128Δ, set2Δ*::NatMX | Stephen McDaniel, unpublished |
| BY4742 | MATα, *his3Δ1 leu2Δ0 lys2Δ0 ura3Δ0* | Open Biosystems, Huntsville, Alabama |
| *bur1Δ BUR1* shuffle (YSB788) | MATα, *ura3-52 leu2Δ trp1Δ63 his3Δ200 lys2Δ202 bur1Δ*::HIS3 *(pRS316-Bur1)* | Keogh et al., 2003 |
| *set2Δbur1Δ BUR1* shuffle (YSB1003) | MATα, *ura3-52* or *3Δ0 leu2Δ1* or *2Δ0 trp1Δ63 his3Δ200* or *3Δ1 lys2Δ202 bur1Δ::*HIS3 *set2Δ*::KanMX *(pRS316-Bur1)* | Keogh et al., 2005 |
| Set2-FRB | MATα ade2-1 trp1-1can1-100 leu2-3, 112 his3-11, 15 ura3 tor1-1 frp1::NAT RPL13A-2XFKBP12::TRP1 Set2-FRB::KanMX6 | this study |
| *set2Δrph1Δ* | MATa, *his3Δ0 leu2Δ0 met15Δ0 ura3Δ0 set2Δ*::HPH *rph1Δ*::KanMX | this study |
| *set2Δjhd1Δ* | MATa, *his3Δ0 leu2Δ0 met15Δ0 ura3Δ0 set2Δ*::HPH *jhd1Δ*::NatMX | this study |
| *set2Δecm5Δ* | MATa, *his3Δ0 leu2Δ0 met15Δ0 ura3Δ0 set2Δ*::HPH *ecm5Δ*::KanMX | this study |
| *set2Δgis1Δ* | MATa, *his3Δ0 leu2Δ0 met15Δ0 ura3Δ0 set2Δ*::HPH *gis1Δ*::KanMX | this study |
| *rph1Δ* | MATa, *his3Δ0 leu2Δ0 met15Δ0 ura3Δ0 rph1Δ*::KanMX | this study |
| *set2Δbar1Δ* | MATa, *his3Δ0 leu2Δ0 met15Δ0 ura3Δ0 set2Δ*::HPH *bar1Δ*::KanMX | this study |
| *set2Δasf1Δ* | MATa, *his3Δ0 leu2Δ0 met15Δ0 ura3Δ0 set2Δ*::NatMX *bar1Δ*::HPH | Stephen McDaniel, unpublished |

#### 2.2 Table S2: Plasmids used in this study and their Addgene deposition IDs.

| Name | Insert | Addgene ID |
|---|---|---|
| mVenus-FLAG-LANS-Set2 | LANS-Set2 | 122002 |
| FLAG-LANS-Set2 | LANS-Set2 | 122003 |

#### 3 Supplemental Movie

**LANS-Set2 light activation and reversion in *Saccharomyces cerevisiae*.** Still images from this video are shown in Figure 1B and quantification of nuclear/cytoplasmic fluorescence intensity change before and during light activation is shown in Figure 1C. Activation is performed on the entire field of view and the appearance and disappearance of a blue circle indicate the time of blue light activation. Scale bar is 3 $\mu$m. (Supplemental_Movie_S1.mov)

#### 4 Supplemental Note

Protein amino acid sequences for mVenus-FLAG-LANS-Set2 and FLAG-LANS-Set2.

mVenus-FLAG-LANS- Set2 sequence (Size: 132.6 kDa):

MKALTVSKGEELFTGVVPILVELDGDVNGHKFSVSGEGEGDATYGKLTLKLICTTGKLP
VPWPTLVTTLGYGLQCFARYPDHMKQHDFFKSAMPEGYVQERTIFFKDDGNYKTRAE
VKFEGDTLVNRIELKGIDFKEDGNILGHKLEYNYNSHNVYITADKQKNGIKANFKIRHNIE
DGGVQLADHYQQNTPIGDGPVLLPDNHYLSYQSKLSKDPNEKRDHMVLLEFVTAAGIT
LGMDELYKGSGSGGRDYKDDDDKSRLALKLAGLDIGLQGSGSGSLATTLERIEKNFVIT
DPRLPDNPIIFASDSFLQLTEYSREEILGRNCRFLQGPETDRATVRKIRDAIDNQTEVTV
QLINYTKSGKKFWNLFHLQPMRDQKGDVQYFIGVQLDGTEHVRDAAEREGVMLIKKTA
ENIDMAAKRVKLDGSGSGSGSRGMSKNQSVSASEDEKEILNNNAEGHKPQRLFDQEP
DLTEEALTKFENLDDCIYANKRIGTFKNNDFMECDCYEEFSDGVNHACDEDSDCINRLT
LIECVNDLCSSCGNDCQNQRFQKKQYAPIAIFKTKHKGYGVRAEQDIEANQFIYEYKGE
VIEEMEFRDRLIDYDQRHFKHFYFMMLQNGEFIDATIKGSLARFCNHSCSPNAYVNKWV
VKDKLRMGIFAQRKILKGEEITFDYNVDRYGAQAQKCYCEEPNCIGFLGGKTQTDAASL
LPQNIADALGVTVSMEKKWLKLKKLSGEPIIKNENENINIEFLQSLEVQPIDSPVDVTKIM
SVLLQQDNKIIASKLLKRLFTIDDDSLRHQAIKLHGYTCFSKMLKLFITEQPQVDGKGNET
EEDDIKFIKGILDFLLELPKTTRNGIESSQIDNVVKTLPAKFPFLKPNCDELLEKWSKFETY
KRITKKDINVAASKMIDLRRVRLPPGWEIIHENGRPLYYNAEQKTKLHYPPSGSSKVFSS
RSNTQVNSPSSSGIPKTPGALDSGSHKLSDEEYEGSGQKRLEYERIALERAKQEELESL
KQKLKLENERKSVLEDIIAEANKQKELQKEEAKKLVEAKEAKRLKRKTVSQSQRLEHNW
NKFFASFVPNLIKKNPQSKQFDHENIKQCAKDIVKILTTKELKKDSSRAPPDDLTKGKRH
KVKEFINSYMDKIILKKKQKKALALSSASTRMSSPPPSTSS

FLAG-LANS- Set2 sequence (Size: 105.7 kDa):

MKALTSGGRDYKDDDDKSRLALKLAGLDIGLQGSGSGSLATTLERIEKNFVITDPRLPD
NPIIFASDSFLQLTEYSREEILGRNCRFLQGPETDRATVRKIRDAIDNQTEVTVQLINYTK
SGKKFWNLFHLQPMRDQKGDVQYFIGVQLDGTEHVRDAAEREGVMLIKKTAENIDMAA
KRVKLDGSGSGSGSRGMSKNQSVSASEDEKEILNNNAEGHKPQRLFDQEPDLTEEAL
TKFENLDDCIYANKRIGTFKNNDFMECDCYEEFSDGVNHACDEDSDCINRLTLIECVND
LCSSCGNDCQNQRFQKKQYAPIAIFKTKHKGYGVRAEQDIEANQFIYEYKGEVIEEMEF
RDRLIDYDQRHFKHFYFMMLQNGEFIDATIKGSLARFCNHSCSPNAYVNKWVVKDKLR
MGIFAQRKILKGEEITFDYNVDRYGAQAQKCYCEEPNCIGFLGGKTQTDAASLLPQNIA
DALGVTVSMEKKWLKLKKLSGEPIIKNENENINIEFLQSLEVQPIDSPVDVTKIMSVLLQQ
DNKIIASKLLKRLFTIDDDSLRHQAIKLHGYTCFSKMLKLFITEQPQVDGKGNETEEDDIK
FIKGILDFLLELPKTTRNGIESSQIDNVVKTLPAKFPFLKPNCDELLEKWSKFETYKRITKK
DINVAASKMIDLRRVRLPPGWEIIHENGRPLYYNAEQKTKLHYPPSGSSKVFSSRSNTQ
VNSPSSSGIPKTPGALDSGSHKLSDEEYEGSGQKRLEYERIALERAKQEELESLKQKLK
LENERKSVLEDIIAEANKQKELQKEEAKKLVEAKEAKRLKRKTVSQSQRLEHNWNKFFA
SFVPNLIKKNPQSKQFDHENIKQCAKDIVKILTTKELKKDSSRAPPDDLTKGKRHKVKEFI
NSYMDKIILKKKQKKALALSSASTRMSSPPPSTSS

#### 5 Supplemental Statistical Methods

##### 5.1 Description of the data

We sought to characterize the methylation dynamics of histone H3 at lysine 36 (H3K36) at a majority of yeast genes in three biological conditions: after light activation of LANS-Set2 (writer add:WA), and after dark inactivation of LANS-Set2 with and without the presence of Rph1, a histone deacetylase (writer loss:WL and writer eraser loss:WEL, respectively). H3K36 trimethyl (H3K36me3) levels were measured through ChIP-seq across four timepoints for each condition (0/20/40/60 min for WA and 0/30/60/90 min for WL and WEL) for 5,355 genes. Seq data were collected and quantified from three sample replicates for each condition across the timepoints. We used two different approaches to process the seq data for analysis. First, within each replicate, H3K36me3 levels for a gene and condition were scaled to the proportion of maximum H3K36me3 level observed (for said replicate, gene, and condition), resulting in a common quantile scale across all replicates, genes, and conditions, which we refer to as "relative H3K36me3 signal" in the text. Second, we took the measurements, previously standardized based on spike-in samples and normalized to adjust for gene length, multiplied them by 1000, and rounded to an integer, resulting in count data (quasi-counts) with a distribution proportional to the starting data.

##### 5.2 General statistical model

Though the H3K36me3 data were large and well-balanced–the vast majority of genes having complete measurements of three replicates from each condition across four timepoints–they possess challenging features from a statistical modeling perspective. The statistical model should accommodate the non-normal distribution of the H3K36me3 data, as well as the time course, *i.e.* longitudinal, nature of observations within a replicate. This broad range of features can be flexibly handled using Bayesian inference (Gelman et al. 2013; Gelman and Hill 2006).

The Stan statistical platform (Carpenter et al. 2017) is one such computational tool for fitting complex Bayesian hierarchical models. We used the brms software package (Bürkner 2017, 2018), which acts as a wrapper of Stan in the R statistical programming language (R Core Team 2019). Let $y_{ijkl}$ be the H3K36me3 measurement for the $i^{\text{th}}$ replicate sample ($i = 1, 2, 3$) at the $j^{\text{th}}$ timepoint ($j = 0, 1, 2, 3$) for the $k^{\text{th}}$ gene ($k = 1, 2, \ldots, 5355$) for $l^{\text{th}}$ condition ($l = 1, 2, 3$). Briefly, we model $y$ with a GLM by defining $E(y) = g^{-1}(\eta)$, where $E(.)$ is the expected value of a random variable, $g^{-1}(.)$ is the inverse link function, and $\eta$ is the linear predictor that relates the outcome to factors of interest.

A Bayesian analysis could in principle simultaneously model all genes and conditions, though in practice, such an approach would likely be computationally infeasible, particularly when considering the need for sufficient sampling in order for the parameter estimates to converge. To avoid these challenges, we instead model more manageable subsets of H3K36me3 data. First, we fit a model of H3K36me3 data for a specific gene and condition. Second, to make more direct comparisons between the writer loss and writer eraser loss conditions, we modeled their data jointly within a gene.

##### 5.3 Zero-one-inflated beta regression model of H3K36me3 quantiles

For the H3K36me3 data transformed to the quantile scale, we initially considered dose response (DR) models (Slob 2002; Wilson et al. 2014). Because the DR model implementations did not easily accommodate covariates or the replicate observations and also fit relatively complex models ($\leq 5$ parameters), we instead modeled the data with a zero-one-inflated beta (ZOIB) distribution with the following parameterization:

$$p(y_i) = \begin{cases} \alpha(1-\gamma) & y_i = 0 \\ \alpha\gamma & y_i = 1 \\ \frac{y_i^{\mu\phi-1}(1-y_i)^{(1-\mu)\phi-1}}{B(\mu\phi,(1-\mu)\phi)} & y_i \notin \{0,1\} \end{cases} \tag{1}$$

$\alpha$ is the zero-one-inflation probability (probability that a zero or one occurs), $\gamma$ is the conditional one-inflation probability (probability that one occurs rather than zerio), $B(.)$ is the beta function (Casella and Berger 2002), and $\phi$ is a positive precision parameter. For the link function, we used the logit: $g(\eta) = \log(\frac{\eta}{1-\eta})$. Zero-one-inflation was necessary because standard beta regression expects $y \in (0, 1)$, meaning it cannot handle values at the boundaries.

The H3K36me3 data scaled to the proportion of the maximum H3K36me3 measured within a replicate (three replicates per gene per condition) results in three values of 1 for each gene and condition pairing. Additionally, zeros may be observed. We used this scale of the data to better standardize ChIP-seq dynamics across cells and better detect consistent patterns compared to the normalized ChIP-seq data. However, it does represent a challenging formulation in the context of a GLMM, and possibly even an abuse of the underlying assumptions of a beta regression model, specifically. The zeros and ones are informative, but would strongly violate the expectations of standard beta regression, resulting in anti-conservative, extreme parameter estimates. To avoid this issue, ZOIB conservatively drops out zeros and ones in terms of parameter estimation, resulting in less extreme estimates for the time effect. Alternatively, we modeled the non-quantile, normalized ChIP-seq data (quasi-counts).

##### 5.4 Zero-inflated negative binomial regression model of H3K36me3 quasi-counts

The normalized ChIP-seq data after being processed and normalized, ranged from 0 to 1.374346. The distribution was consistent in shape with common count distributions, *e.g.* Poisson and negative binomial, exemplified by being non-negative with a right skew. We calculated quasi-counts as $y_i^{\text{quasi}} = \text{round}(y_i \times 1000)$. Though this transformation is artificial, it produces a new distribution that is proportional to the original and consistent with count distributions.

For the quasi-count scale, we modeled the data with a zero-inflated negative binomial (ZINB) with the following parameterization:

$$p(y_i) = \begin{cases} \binom{y_i+\phi-1}{y_i}(\frac{\mu}{\mu+\phi})^{y_i}(\frac{\phi}{\mu+\phi})^{\phi} & y_i > 0 \\ \xi + (1-\xi)\text{NB}(y_i = 0) & y_i = 0 \end{cases} \tag{2}$$

$\xi$ is the zero-inflation probability, NB$(.)$ is the non-zero-inflated negative binomial probability mass function, and $\phi$ is a positive precision parameter. As $\phi \to \infty$, the negative binomial distribution converges to the Poisson distribution. The ZINB distribution estimates the zero observations as a mixture of true zeros, expected by the NB distribution, with an additional component from drop-outs, resulting in the zeros having less influence on the parameter estimates.

The inference on the H3K36me3 time rate dynamics based on either ZOIB and ZINB should be consistent with each other. The quantile data scale is more standardized across genes and conditions, but also overly conservative when modeled by ZOIB regression, due to essentially removing the effect of zeros and ones on parameter inference. By contrast, the quasi-count scale is more artificial and disparate across genes and conditions, but more closely represented the raw data. It also makes more complete use of the data–by not excluding the maximum values which was transformed to one in the quantile data–for parameter estimation.

##### 5.5 Time models

###### *5.5.1 Continuous time model*

We modeled the gain or loss of H3K36me3 with a continuous time variable in the linear predictor of the model,

$$\eta_{ij} = \mu + u_i + (\beta_{\text{time}} + v_i)x_{ij}, \tag{3}$$

where $\mu$ is a shared intercept term, $x_{ij}$ is the $j^{\text{th}}$ timepoint (in minutes) for the $i^{\text{th}}$ sample, and $\beta_{\text{time}}$ is the change rate with time. $u_i$ and $v_i$ are random, or group-level (Gelman and Hill 2006), effects that account for the longitudinal nature of the data, and modeled with the following priors: $\text{N}(0, \tau_u^2)$ and $\text{N}(0, \tau_v^2)$, respectively. For all genes and conditions, we recorded the posterior mean ($\widehat{\beta}_{\text{time},kl}$) as a point estimate for the change in H3K36me3 (proportion or quasi-count) with time for gene $k$ and condition $l$, the standard error on the estimate, and the 95% Credible Interval (CrI), which we used to define "confident" genes (CrI that do not cover 0) for a given condition. $\widehat{\beta}_{\text{time},kl}$ were plotted and correlated with additional covariates of interests in order to identify potential relationships with other factors of interest.

##### 5.5.2 *Testing for non-zero effect of condition with time*

To formally test for a non-zero effect of condition with time is infeasible because it would require the joint estimation across all genes and conditions in a Bayesian context. Further, GLMMs are challenging models to fit, and not amenable to reliable likelihood-based inference, given the number of genes observed here. As an alternative, we fit a second model from the posterior mean $\widehat{\beta}_{\text{time},kl}$, estimated for gene $k$ and condition $l$, described above. These parameters were modeled as normally distributed, which can be viewed as a latent variable that we model in a second regression as:

$$\beta_{\text{time},kl} = u_k + \delta_{\text{WA}} I\{l = \text{WA}\} + \delta_{\text{WL}} I\{l = \text{WL}\} + \delta_{\text{WEL}} I\{l = \text{WEL}\} + \varepsilon_{kl}, \tag{4}$$

where $u_k$ is a gene-specific random effect, modeled as $\text{N}(0, \tau_u^2)$, $\delta_{\text{WA}}$, $\delta_{\text{WL}}$, and $\delta_{\text{WEL}}$ are the condition-specific effects on the previously measured continuous time effect, modeled as fixed effects, and $\varepsilon_{kl}$ is a noise term, distributed according to $\text{N}(0, \frac{\sigma^2}{w_{ij}})$ with $\sigma^2$ representing the noise variance and $w_{kl}$ is a weight specific to gene $k$ and condition $l$. For the weight, we used $1/\text{SE}(\beta_{\text{time},kl})$ from the GLMM model, effectively down weighting the influence of effect estimates with large standard error. $I\{A\}$ represents the indicator function, returning 1 if the conditional statement A is satisfied and 0 if not.

Given that the effects can be modeled with a normal distribution, representing a linear mixed effect model (LMM) with weights, we used the R package lme4 (Bates et al. 2015). Through ANOVA (Venables and Ripley 2010) with maximum likelihood estimates, we compared the model in Equation 4 to a null model with an intercept and no condition fixed effects, resulting in an ANOVA $p$-value $< 2.2 \times 10^{-16}$. Using the R package emmeans (Lenth 2019), we performed Tukey *post hoc* tests of pairwise differences (Venables and Ripley 2010) between the conditions, which were all found to be significant (Tukey $p$-values $< 0.0001$). Notably, the rate of H3K36me3 loss was greater for WEL compared to WL.

##### 5.5.3 *Categorical time model*

WL and WEL had similar H3K36me3 dynamics with time. To more directly compare their dynamics, we simultaneously analyzed WL and WEL data per gene within a categorical time model, allowing for greater flexibility and non-linearity with respect to time:

$$\begin{aligned}\eta_{ijl} = &\underbrace{\mu + u_i}_{\text{WL time0}} + \underbrace{(\omega_{\text{WEL}} + w_i) I\{l = \text{WEL}\}}_{\text{WEL time0}} \\ &+ \underbrace{\sum_{p=1}^{3} (\beta^p_{\text{timepoint}} + v^p_i) I\{j = p\}}_{\text{WL timepoint } p} \\ &+ \underbrace{\sum_{p=1}^{3} (\delta^p_{\text{timepoint}} + z^p_i) I\{j = p, l = \text{WEL}\}}_{\text{WEL timepoint } p},\end{aligned} \tag{5}$$

where $\mu$ is the intercept representing, representing WL at time0, $\omega_{\text{WL}}$ is the deviation of WEL from WL, $\beta^p_{\text{timepoint}}$ is the effect comparing timepoint $p$ to 0 for WL, and $\delta^q_{\text{timepoint}}$ represents effect for timepoint $p$ for WEL compared to WL. $u_i$, $w_i$, $v^p_i$, $z^p_i$ are group-level effects that model the correlations within replicates, and have the following prior distributions: $\text{N}(0, \tau^2_u)$, $\text{N}(0, \tau^2_w)$, $\text{N}(0, \tau^2_{v,p})$, and $\text{N}(0, \tau^2_{z,p})$, respectively. Similar to the continuous time model, we recorded posterior means and 95% CrI for $\delta^p_{\text{timepoint}}$. We fit the categorial time model with the quantile data and ZOIB, though it could be used with the quasi-counts and ZINB as well.

*5.5.4 Interpreting estimated effects*

The interpretation of the regression coefficients will depend on whether ZOIB or ZINB was used. For ZOIB, effects represent log odds ratio (OR) for change in proportion H3K36me3 (relative to the sample maximum) per minute. For ZOIB, the effects are log change in quasi-counts per minute. We emphasize that these data were highly derived, resulting in some reduction in the interpretability or tangibility of their estimates. Regardless of data scale, positive effects represent increasing levels of trimethyl marks and negative effects represent decreasing marks. We view large scale trends across genes and/or conditions as meaningful.

##### 5.6 Model specification, sampling, and convergence

A Bayesian model requires the specification of prior distributions for the various parameters. We used the default settings from brms. Briefly, for fixed, or population-level, effects, an improper flat prior over the reals was used. Group-level effects are modeled as normal variables with standard deviation parameter. These are modeled with half Student-$t$ distributions with 3 degrees of freedom and a minimal scale parameter of 10 (Gelman and Hill 2006), which brms potentially increases based on the data to insure that the prior is minimally informative. The LKJ-correlation prior is used to model the correlations between the group-level effects on the same grouping factor (Lewandowski et al. 2009).

Bayesian inferences involves random sampling from the joint posterior of the model parameters. For each Stan model, we ran four Monte Carlo Markov Chains (MCMC), each with 2,000 iterations of warm-up and sampling. Initial values for parameters for each chain were randomly generated within Stan. The the adaptive delta parameter, necessary for the No-U-turn Sampler (Hoffman and Gelman 2014) used by Stan, was set to 0.8, the default used by brms.

There are various diagnostics for the MCMC samples that can be used to determining whether the model is converging and performing well. We used the split-$\widehat{R}$ statistic (Gelman et al. 2013), a ratio of the average variance within chains to the variance with the pooled chains. A split-$\widehat{R}$ value close to 1 means the variances within each chain are similar, and thus the model is likely mixing well and converging. Guidelines from the Stan development team state that models with split-$\widehat{R} > 1.1$ have not converged and should not be used for inference. To ensure replicable results and declare convergence and ensure reproducible results, we set the seed and ran all models in Stan (through brms) for a given gene. We then checked that split-$\widehat{R} < 1.1$ for all parameters. If this was satisfied, we declared convergence and recorded the seed as well as posterior means and CrIs for the parameters. If any split-$\widehat{R} > 1.1$, convergence failed, and we set a new seed and repeated the process. We capped that number of repeats at 10. If convergence is not met after 10 iterations, we declare modeling to have failed for those genes, likely due to oddly distributed and noisy data.

R Core Team. RSoftware2019, 2019.

Wout Slob. Dose-Response Modeling of Continuous Endpoints. *Toxicological Sciences*, 66(2):298–312, apr 2002. ISSN 10960929. doi: 10.1093/toxsci/66.2.298. URL https://academic.oup.com/toxsci/article-lookup/doi/10.1093/toxsci/66.2.298.

W. N. Venables and B. D. Ripley. *Modern Applied Statistics with S*. Springer Publishing Company, Incorporated, 2010. ISBN 1441930086, 9781441930088.

Ander Wilson, David M. Reif, and Brian J. Reich. Hierarchical dose-response modeling for high-throughput toxicity screening of environmental chemicals. *Biometrics*, 70(1):237–246, mar 2014. ISSN 0006341X. doi: 10.1111/biom.12114. URL http://doi.wiley.com/10.1111/biom.12114.
